# Unexpected transformations during pyrroloiminoquinone biosynthesis

**DOI:** 10.1101/2024.03.12.584671

**Authors:** Josseline Ramos Figueroa, Lingyang Zhu, Wilfred A. van der Donk

## Abstract

Pyrroloiminoquinone containing natural products have long been known for their biological activities. They are derived from tryptophan, but their biosynthetic pathways have remained elusive. Studies on the biosynthetic gene cluster (BGC) that produces the ammosamides revealed that the first step is attachment of Trp to the C-terminus of a scaffold peptide in an ATP and tRNA dependent manner catalyzed by a PEptide Amino-acyl tRNA ligase (PEARL). The indole of the Trp is then oxidized to a hydroxyquinone. We previously proposed a chemically plausible and streamlined pathway for converting this intermediate to the ammosamides using additional enzymes encoded in the BGC. In this study, we report the activity of four additional enzymes that show that the proposed pathway is incorrect and that Nature’s route towards pyrroloiminoquinones is much more complicated. We demonstrate that, surprisingly, the amino groups in pyrroloiminoquinones are derived from three different sources, glycine, asparagine, and leucine, all introduced in a tRNA dependent manner. We also show that an FAD-dependent putative glycine oxidase is required for the process that incorporates the nitrogens from glycine and leucine, and that a quinone reductase is required for the incorporation of the asparagine. Additionally, we provide the first insights into the evolutionary origin of the PEARLs as well as related enzymes such as the glutamyl-tRNA dependent dehydratases involved in the biosynthesis of lanthipeptides and thiopeptides. These enzymes appear to all have descended from the ATP-GRASP protein family.

## Introduction

Peptide aminoacyl-tRNA ligases (PEARLs) catalyze the appendage of amino acids to the C-terminus of scaffold peptides in an ATP and aminoacyl-tRNA dependent manner.^1^ ATP is used to phosphorylate the C-terminus of their peptide substrates to form an acylphosphate intermediate, which is attacked by an aminoacyl-tRNA to generate a new peptide bond (Figure 1A).^2^ Subsequent hydrolysis of the tRNA results in the net addition of an amino acid to the peptide. These enzymes have been shown to add a wide variety of amino acids to the C-termini of their peptide substrates. The first examples involved the installation of cysteine residues to a precursor peptide en route to the biosynthesis of 3-thiaglutamate and related molecules.^1–3^ More recently, PEARL enzymes that are involved in the biosynthesis of pyrroloiminoquinone-derived natural products were characterized.^4^ These latter studies investigated two biosynthetic gene clusters (BGCs, Figure 1B) that share a highly similar precursor peptide, the ammosamide BGC (*amm*) and a gene cluster in *Bacillus halodurans* C-125 (*bha*). PEARL enzymes from these BGCs were shown to append a Trp moiety to a precursor peptide that serves as the core of the pyrroloiminoquinones (Figure 1C). Trihydroxylation of the added Trp residue is catalyzed by an FMN-dependent oxidoreductase (BhaC_1_ or AmmC_1_). While the initial product of the flavoprotein is in the hydroquinone state, oxidation to the quinone form is needed to activate this biosynthetic intermediate for the next PEARL-catalyzed reaction. BhaB_5_ was shown to modify the vinylogous carboxylate at C5 in the oxidized Trp by attaching a glycine residue transferred from Gly-tRNA^Gly^ (Figure 1C).^4^

In the reaction catalyzed by BhaB_5_, a complex mixture of products was obtained. Based on LC-MS and NMR characterization, they were assigned as the tryptophanyl derived glycine-quinone adduct (**1**), a decarboxylated product (**2**, or its methylamine tautomer), and an aminoquinone (**3**) (Figure 1C). We proposed that the glycine-quinone adduct **1** was an off-pathway intermediate formed by dissociation from the enzyme active site. Indeed, the analogous reaction catalyzed by AmmB_3_ in *E. coli* resulted only in production of a peptide similar to **2** derived from the AmmA substrate. We hypothesized that the amino quinone product **3** requires first oxidation of **2** by an oxidase present in *E. coli*, resulting in hydrolysis to release formaldehyde and formation of the amino group that is present in the final product ammosamide. We anticipated that in the producing organism this oxidation would be catalyzed by one of the remaining enzymes in the BGC. We also suggested that the two remaining PEARLs in the ammosamide BGC (AmmB_1_ and AmmB_4_) would utilize Gly-tRNA^Gly^ to introduce the additional two amino groups in the final product (Figure 1D, Figure S1). We show herein that none of these hypotheses are correct and that the biosynthesis of ammosamide and related molecules is more complex than anticipated.

With the aim to find the next enzymatic steps in the biosynthetic pathways encoded by the two gene clusters described above, we investigated the activity of four additional enzymes that are encoded in the BGCs shown in Figure 1B. We show that the aforementioned aminoquinone **3** and the decarboxylated product (**2**) are non-enzymatic decomposition products of the unstable glycine-quinone adduct **1**. In the actual biosynthetic pathway, the latter is modified by a predicted glycine oxidase encoded in both BGCs forming the aminoquinone product through glyoxylate release. Investigation of the remaining unassigned PEARL enzymes of both clusters revealed new PEARL-catalyzed transformations. In the *B. halodurans* C-125 pathway, after formation of aminoquinone **3**, an asparagine is installed to the C-terminus of the modified peptide in an Asn-tRNA^Asn^ dependent fashion. In the ammosamide pathway, after aminoquinone formation, leucine (and not glycine) was added to the aminoquinone providing the second amino group present in the ammosamide core. These findings show that the biosynthetic logic that led to the evolution of the pathway towards ammosamide and related products is complex. Through bioinformatic and structural analysis, we provide new insights into the potential evolutionary origin of PEARLs, and by extension glutamyl-tRNA dependent lanthipeptide and thiopeptide dehydratases.^5^

**Figure 1.**
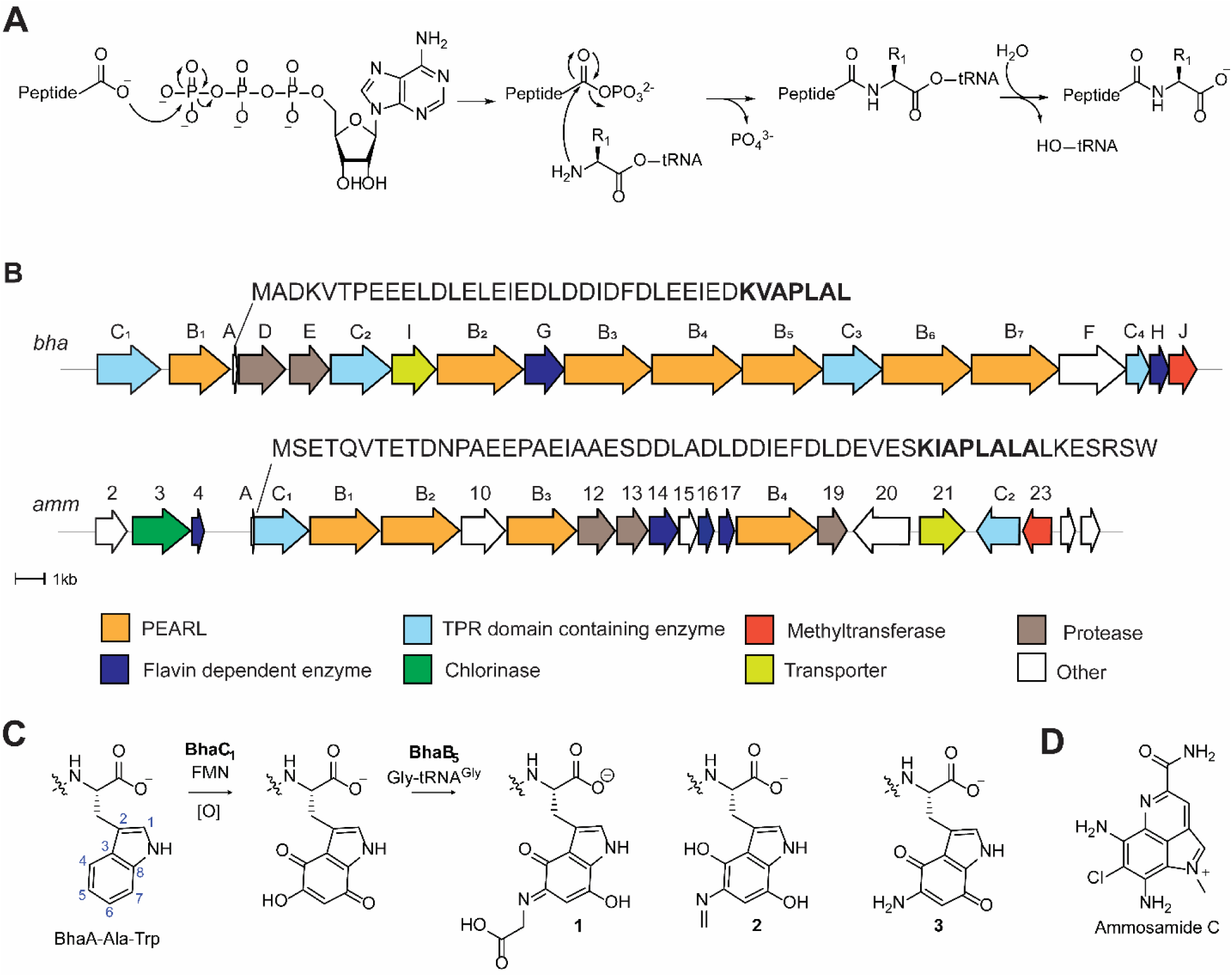
Biosynthesis of pyrroloiminoquinones and related compounds. (A) General reaction scheme for peptide amino-acyl tRNA ligases (PEARLs). (B) Two related BGCs that are thought to result in pyrroloiminoquinone products. The *amm* BGC has been demonstrated to generate ammosamide C; the product of the *bha* BGC is currently not known. (C) After addition of an Ala and Trp at the C-terminus of the precursor peptide BhaA, BhaC_1_ oxidizes the indole of the Trp, and the PEARL BhaB_5_ adds Gly from Gly-tRNA^Gly^ resulting in three observed products. Similar reactions are catalyzed by AmmC_1_ and AmmB_3_. (D) Structure of ammosamide C.

## Results and discussion

### The glycine oxidase BhaG oxidizes the glycine-quinone adduct 1

To uncover the next biosynthetic step in the *bha* pathway, the peptide BhaA-Ala-Trp was heterologously co-expressed in the presence of the previously characterized modifying enzymes (BhaC_1_ and BhaB_5_; Figure 1C)^4,6^ as well as iterative inclusion of each of the remaining enzymes encoded in the *bha* cluster (Figure 1B). After peptide purification and desalting, the samples were submitted for analysis by matrix-assisted laser desorption/ionization time-of-flight (MALDI-TOF) mass spectrometry (MS). Out of all individual co-expression experiments, only the inclusion of a putative glycine oxidase, BhaG (UniProt Q9KB87), resulted in production of a new peak with a decreased mass of 57 Da from the glycine-adduct **1** (Figure 2A). We confirmed that the modification occurred at the C-terminal Trp using high-resolution MS/MS on a trypsin generated fragment as shown in Figure 2C.

Analysis by liquid chromatography (LC) coupled with electrospray ionization (ESI) mass spectrometry showed that peptide **1** was the major product in the absence of BhaG (Figure S2). This result suggested that **1** was not an off-pathway product as previously hypothesized, but the actual product of the reaction catalyzed by BhaB_5_. To confirm that the glycine adduct **1** is converted by BhaG to the aminoquinone **3**, the enzyme was purified and used for *in vitro* reactions. His_6_-BhaG co-purified with FAD (Figure S3) in agreement with other glycine oxidases.^7^ Compounds **1**, **2**, and **3** co-elute under our purification protocols and could not be separated. Therefore, the mixture was used for *in vitro* studies, which contained **1** as the major component (Figure S2). *In vitro* reconstitution of BhaG activity showed that the glycine-adduct **1** was converted into the amino quinone product **3** (Figure S4). The enzymatic conversion was expected to involve hydride transfer from the α-carbon of the glycine moiety to FAD, then hydration of the resulting imine to generate a tetrahedral intermediate that eliminates glyoxylate and forms **3** (Figure S5A). Hence, we investigated whether glyoxylate was formed over the course of the reaction by derivatization of the reaction products using phenylhydrazine (Figure S5B). As expected, the phenylhydrazone adduct of glyoxylate was formed as shown by co-injection with a standard mixture obtained from glyoxylic acid. Thus, the Gly adduct **1** is not converted to **3** by decarboxylation, oxidation, and hydrolysis to provide formaldehyde as previously proposed,^4^ but instead by enzymatic oxidation by a glycine oxidase that is conserved in both BGCs.

**Figure 2.**
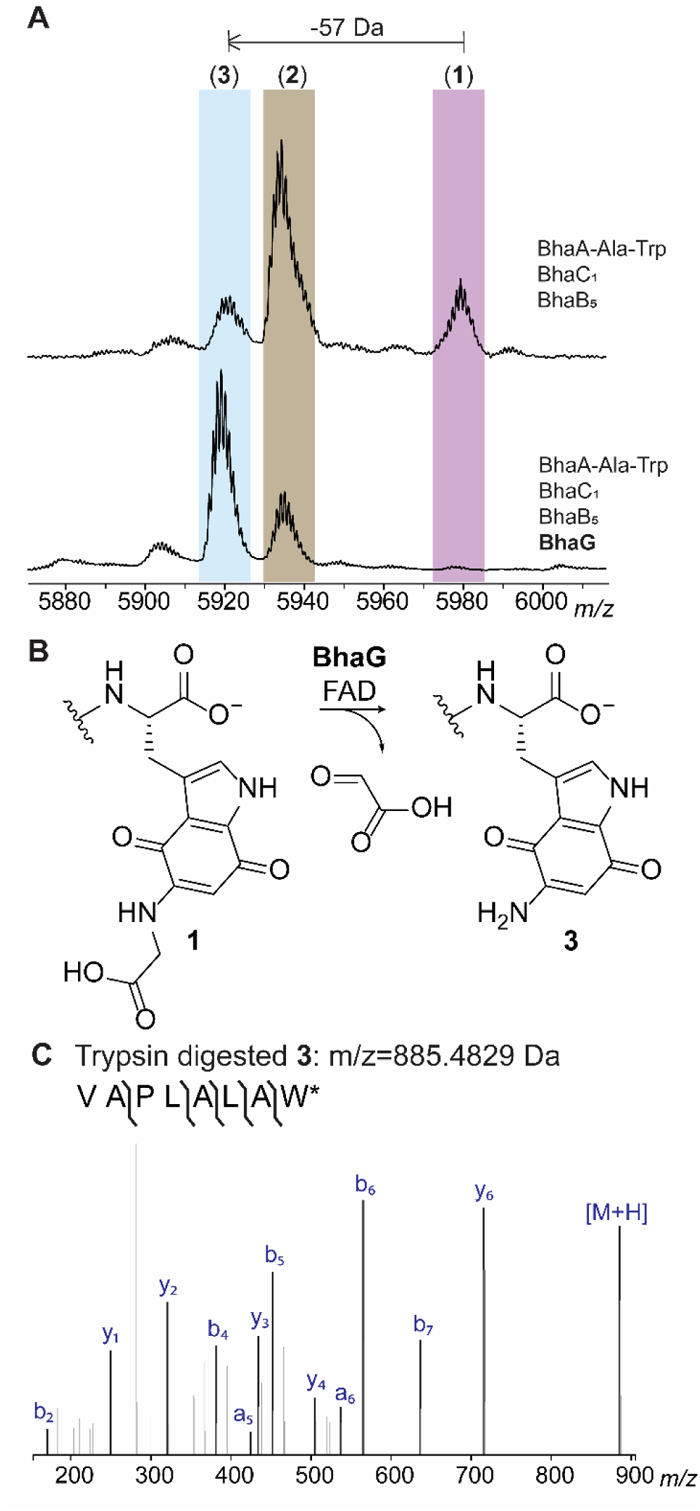
Activity of BhaG. (A) Co-expression of Bha-Ala-Trp with BhaC_1_ and BhaB_5_ in *E. coli* and subsequent MALDI-TOF MS analysis results in detection of the glycine adduct **1**, the imine **2**, and the aminoquinone **3**. Inclusion of BhaG in the co-expression experiment leads to disappearance of the glycine adduct **1**, strong reduction of the peak of peptide **2** and formation of predominately **3**. Differences in ionization efficiency in MALDI-TOF MS underestimate the relative amount of product **3** (see LC-MS data in Figure S4). (B) Reaction catalyzed by BhaG. (C) ESI-MS/MS analysis of the C-terminal peptide of trypsin-digested product **3** formed in vitro by BhaG (calculated m/z = 885.4829, observed m/z = 885.4827).

### BhaB4 is an Asn-tRNA^Asn^ dependent PEARL

In previous experiments we were unable to identify the next PEARL to act in the pathway, which may be explained by the need to include BhaG to efficiently generate peptide **3**. Thus, we investigated whether this new information might allow the identification of the next biosynthetic step. We co-expressed BhaA-Ala-Trp, BhaC_1_, BhaB_5_ and BhaG and in turn added genes encoding each of the remaining PEARL enzymes to the expression system. As shown in Figure 3A, BhaB_4_ (UniProt Q9KB87) resulted in an increase in the peptide mass by 114 Da whereas the addition of the other PEARLs did not yield any observable mass shift. High resolution MS/MS analysis showed that the increased mass was generated by asparagine condensation to the C-terminal carboxylate of the modified tryptophan (Figure 3B,C). To confirm that asparagine was the amino acid added by BhaB_4_ as well as the site of the new amide bond formation, NMR analysis of the peptide was performed. Confirmation that the Asn residue was added to the C-terminus to produce peptide **4** was provided by a characteristic NOESY peak from the NH proton (δ 7.95 ppm) of the newly added Asn to the beta protons (δ 3.10 and 2.87 ppm) of the modified Trp (Figure S6A).

Next, we investigated whether the activity of BhaB_4_ could be recapitulated *in vitro* using purified aminoquinone intermediate **3**. Initial attempts to detect the desired mass shift using tRNA^Asn^, Asn-tRNA^Asn^ transferase from *E. coli* (AsnRS), BhaB_4_, ATP, **3** and Asn did not result in product formation. However, inclusion of an aliquot of *E. coli* cell free extract did result in formation of the +114 Da product (Figure S7). We hypothesized that an enzyme in the *E. coli* extract was modifying the redox state of the aminoquinone **3**, which is obtained in the oxidized form in our purification protocols. This hypothesis was further fueled by the presence of a putative NADPH-dependent FMN-quinone reductase in the *bha* cluster (BhaH, UniProt Q9KB78, Figure 1A). Thus, we investigated whether the need for *E. coli* cell free extract could be obviated by inclusion of BhaH.

**Figure 3.**
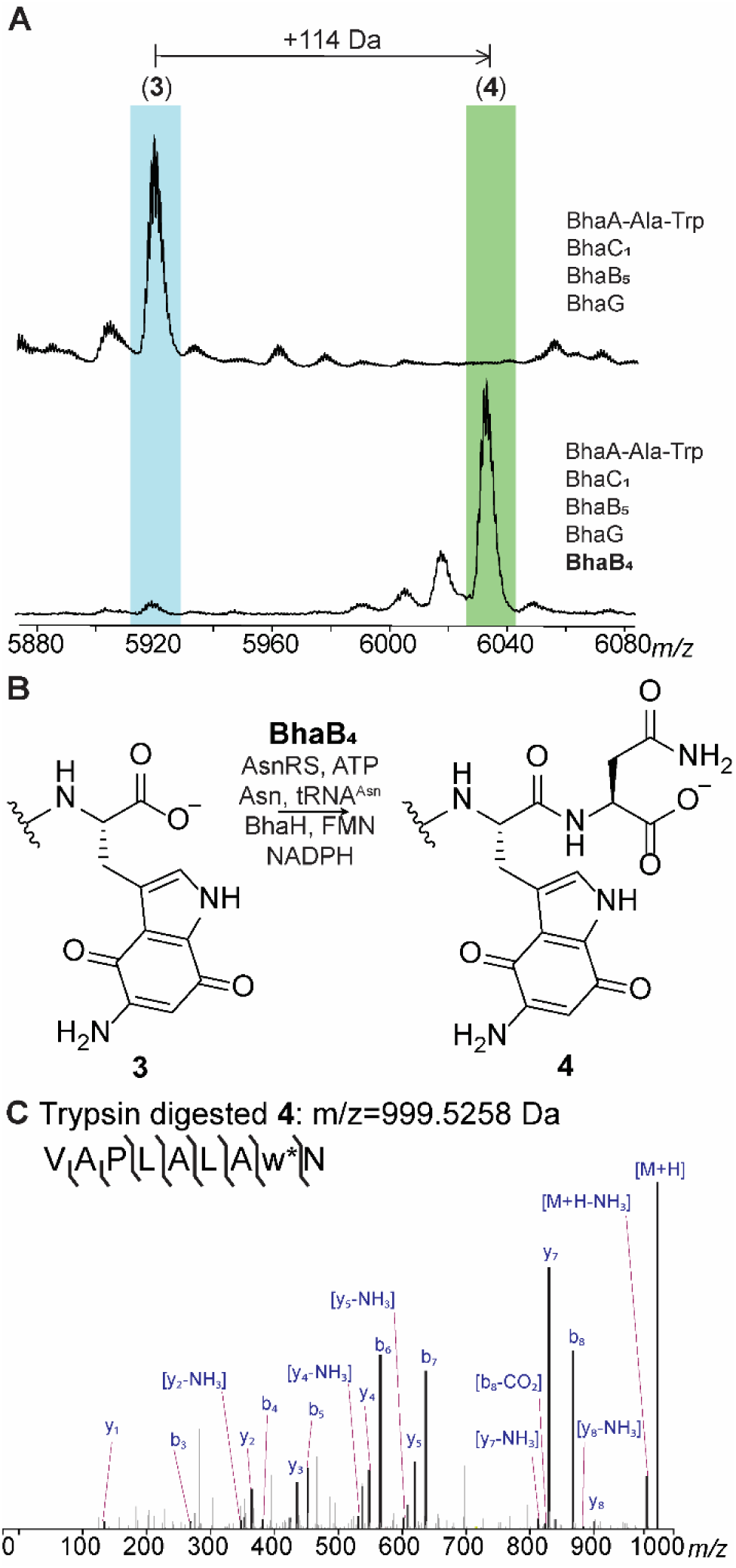
BhaB_4_ adds Asn to the C-terminus of intermediate **3**. (A) Inclusion of BhaB_4_ in the co-expression system that produces **3** results in the formation of a new product that is increased in mass by 114 Da. (B) The reaction catalyzed by BhaB_4_. (C) ESI-MS/MS analysis of the C-terminal fragment upon trypsin digestion of peptide **4** (calculated m/z = 999.5258, observed m/z = 999.5282; Figure S7B).

We expressed BhaH as an N-terminal His_6_-tagged fusion protein that co-purified with FMN. The purified enzyme reduced the standard substrate menadione in the presence of NADPH (Figure S8). We next investigated whether the aminoquinone **3** would be a substrate for Asn conjugation by BhaB_4_ in the presence of BhaH. We previously observed rapid oxidation of the hydroquinones in the *bha* pathway under ambient conditions,^4^ and because BhaH was anticipated to generate such a hydroquinone from intermediate **3** we performed the asparagine condensation reaction catalyzed by BhaB_4_ under anaerobic conditions in the presence of BhaH and NADPH. Indeed, we observed complete formation of the +114 product as shown by MALDI-TOF MS and high-resolution MS/MS analysis whereas no product formation was observed when BhaH or NADPH was omitted from the reaction mixture (Figures S7A, S8C). Having identified conditions for full conversion, we then used the anaerobic conditions with isotopically labeled L-^15^N_2_-Asn for the *in vitro* assay with BhaB_4_ and BhaH. Using LC-MS analysis, the isotopically labeled asparagine was incorporated in the product (M+2 Da; Figure S9). Thus, the substrate for BhaB_4_ is the likely the reduced hydroquinone form of **3**, which is then condensed with Asn at the C-terminal carboxylate in an Asn-tRNA^Asn^ and ATP dependent manner (Figure 4A). This product oxidizes upon exposure to oxygen during sample preparation for high-resolution MS/MS analysis providing peptide **4**. The reason why peptide **3** might need to be reduced by BhaH for BhaB_4_ activity is not entirely clear. One possibility is that it prevents reaction at the vinylogous carboxylate that is present at C7 in the imine tautomer of **3**; such a vinylogous carboxylate at C5 is the site of condensation by BhaB_5_ (Figure 1B).^4^

### A different glycine-adduct is obtained in the *amm* pathway than in the *bha* pathway

In a previous study we demonstrated using NMR spectroscopy that in the *bha* pathway, a glycine moiety is added by BhaB_5_ to the Trp core at position 5 of the indole moiety to generate **1** (Figure 1C).^4^ In the same study it was also shown by MS that AmmB_3_ adds a Gly to the oxidized indole of a C-terminal Trp in its substrate (AmmA*-Trp), but the product was not characterized by NMR spectroscopy. Since the final product of the *amm* BGC is known (ammosamide C, Fig. 1D), two amine groups need to be installed onto the indole. Although it appeared most likely that AmmB_3_ would also add Gly to C5, like BhaB_5_, we could not rule out that tautomerization involving positions 5 and 7 of the quinone in the substrate would result in activation of the latter for the condensation step.^4^ Hence, in the current work we co-expressed His_6_-AmmA*-Trp, AmmC_1_, and AmmB_3_ in *E. coli* in a large scale (over 30 L) and purified the product peptide **5** for NMR characterization. After trypsin digestion, the C-terminal fragment containing the modified IAPLALAw (w = modified Trp) sequence was obtained and analyzed by multi-dimensional NMR spectroscopy. The data revealed that the site of glycine addition by AmmB_3_ is different from that observed for BhaB_5_. ^1^H-^13^C HMBC revealed correlations from the NH proton of the glycine adduct (δ 7.82 ppm) to the indole carbon at C8 (δ 131.90 ppm) indicating that the glycine addition occurred at C7 (intermediate **5**, Figure 4B). Nuclear Overhauser Effect Spectroscopy (NOESY) data confirmed this assignment by the observation of NOE cross peaks from the NH proton (δ 7.82 ppm) of the appended glycine to the NH proton (δ 11.92 ppm) of the indole moiety as well as from the CH_2_ protons (δ 4.10 ppm) of the appended glycine to the indole CH proton at C6 (δ 5.00 ppm) (Figure S10). As observed previously for the C5 adduct in the *bha* pathway,^4^ the glycine adduct at C7 was unstable and decomposed over time during our NMR measurements.

**Figure 4.**
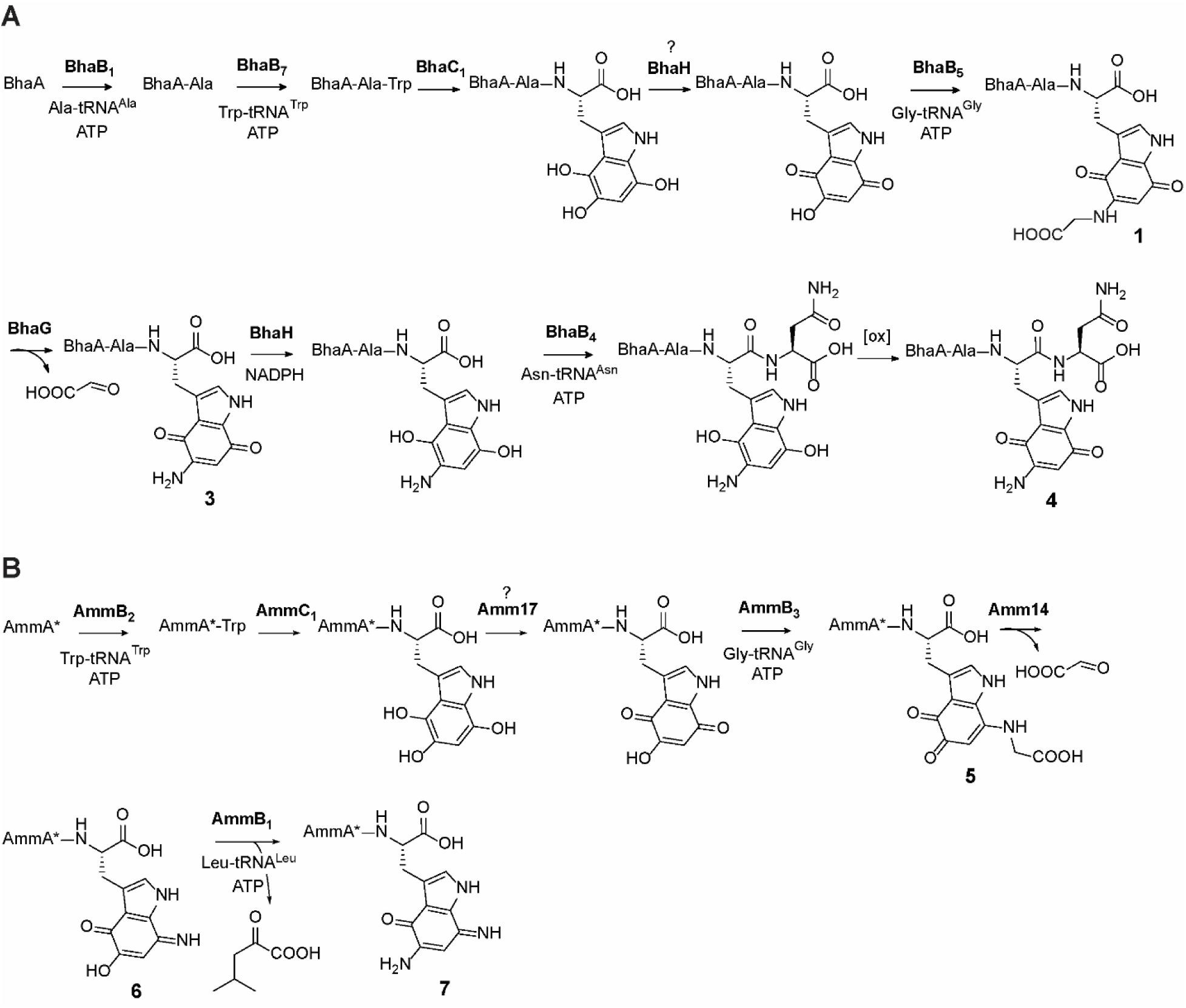
Updated functions of biosynthetic enzymes from the *bha* and *amm* BGCs. (A) Pathway encoded by the *bha* BGC based on the data in this and previous studies. (B) Pathway towards ammosamide C encoded by the *amm* BGC based on the data in this and previous studies.

### Further insights into the flavin-dependent indole trihydroxylase BhaC_1_

The observation that AmmB_3_ introduces an amino group from Gly-tRNA at C7 of the hydroxyquinone to give **5** whereas BhaB_5_ catalyzes the same reaction at C5 to give **1** (Figure 4) may also explain why in the absence of the glycine oxidases AmmG/BhaG the major observed products formed by co-expression of the respective enzymes in *E. coli* are different. For the Bha pathway, the major product is the glycine adduct **1** at C5 (Figure S2) but for the Amm pathway it is the imine **2’** at C7 (or its tautomer, Figure S10E).^4^ Apparently, the decarboxylation of the glycine adduct at C7 is easier than at C5. The different reactions catalyzed by BhaB_5_ and AmmB_3_ also may explain one additional observation. When we co-expressed His_6_-BhaA-Ala-Trp with BhaC_1_ and BhaB_5_, we consistently observed a new band by sodium dodecylsulfate polyacrylamide gel electrophoresis that runs at slightly higher molecular weight than BhaC_1_ (Figure 5A). Such a band was not observed when BhaB_5_ was omitted nor was it observed for the equivalent experiment with AmmA*-Trp, AmmC_1_ and AmmB_3_. The protein that is associated with this band contains a His-tag as shown by Western blot analysis (Figure 5A). We isolated the band from the gel, digested the protein with LysC, and analyzed the digest by mass spectrometry. Fragments corresponding to sequences of both BhaC_1_ and BhaA-Ala-Trp were observed suggesting the band corresponds to a covalently crosslinked protein. Indeed, a fragment was observed that links a peptide spanning residues 382-432 of BhaC_1_ to the C-terminus of modified BhaA-Trp with tandem MS analysis suggesting the linkage is between Lys391 and the aminoquinone of modified BhaA (Figure 5B, Figure S11). The reaction catalyzed by BhaC_1_, trihydroxylation of the indole of Trp, is unique in biochemistry and bioinformatics did not predict this tetratrico repeat domain-containing protein to be a flavoprotein.^4,6^ Because the protein does not contain any canonical flavin binding sites, we previously used AlphaFold Multimer^8^ to predict the interaction between the substrate peptide (BhaA-AlaTrp) and BhaC_1_,^6^ and thereby the putative position of the active site of BhaC_1_. The observed covalent interaction between Lys391 of BhaC_1_ and the oxidized indole of BhaA-AlaTrp provides further support for the postulated active site position as Lys391 was predicted to interact with the C-terminal carboxylate of the BhaA-AlaTrp peptide (Figure 5C). The high reactivity of the BhaB_5_ product was reported previously and apparently it results in crosslinking to BhaC_1_ when the amino group is at C5, but not crosslinking to AmmC_1_ when the amino group is at C7. Importantly, the observed adduct is consistent with the mechanism of substrate engagement predicted by AlphaFold Multimer.

**Figure 5.**
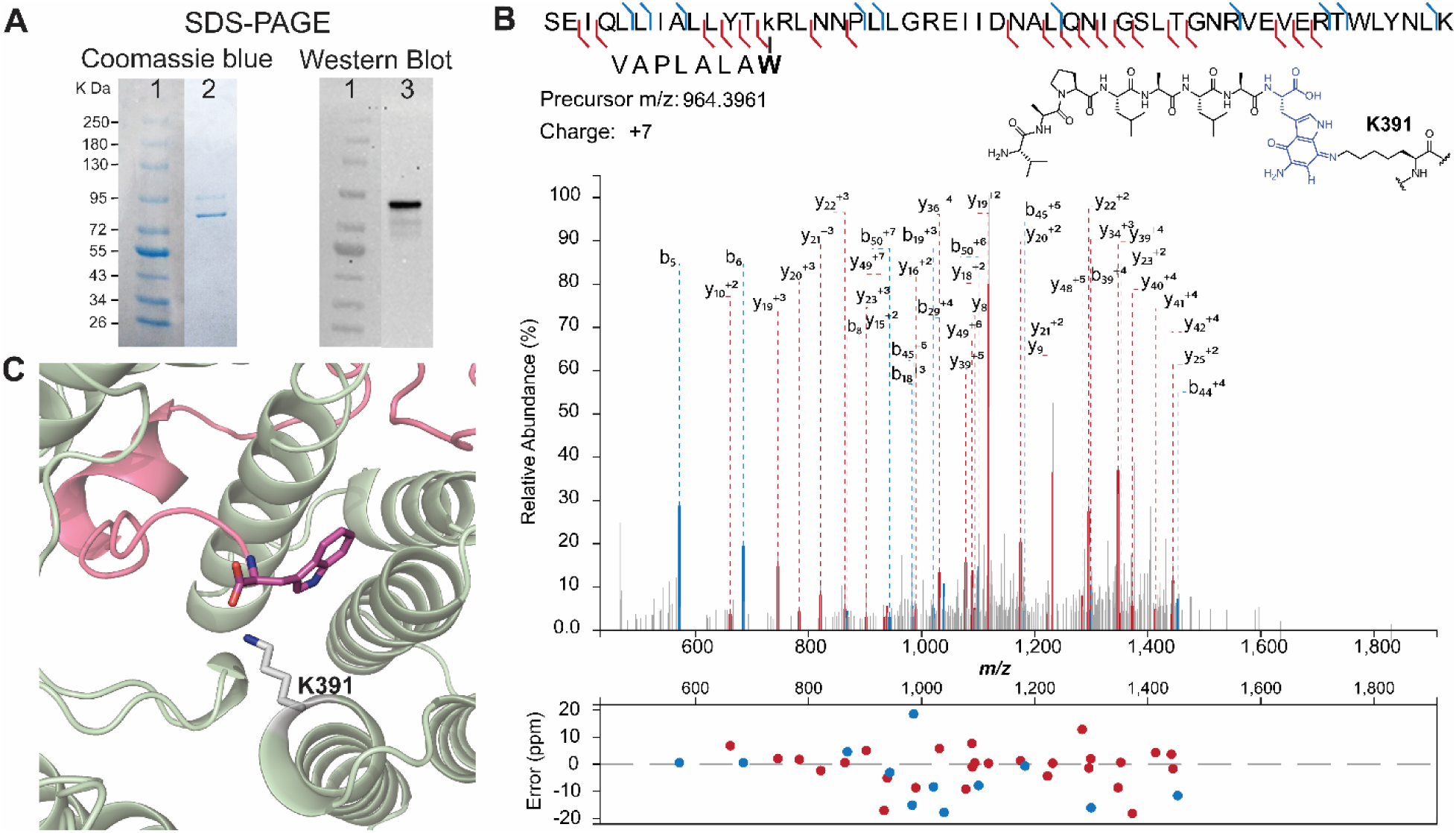
(A) SDS-PAGE analysis of the co-expression product obtained from non-His-tagged BhaC_1_ and non-His-tagged BhaB_5_ co-expressed with His-tagged BhaA-AW. The Coomassie blue stained gel (lane 2) shows two high molecular weight bands around ∼90 kDa and ∼80 kDa. Western Blot analysis (lane 3) showed that the top band contains a histidine tag, likely corresponding to a covalent crosslink of intermediate **3** to BhaC_1_. (B) LC-MSMS data on a LysC-digest of the upper band. The fragment shown consists of a LysC-fragment of BhaC_1_ crosslinked to the C-terminal fragment of modified Bha-AW. The crosslink is drawn at C7 of the indole, but it could also be at C4. For additional tandem MS data supporting the assignment, see Figure S11. (C) Alphafold multimer prediction of the binding of BhaC1 (colored sage) to BhaA_AW (pink) positioning the C-terminal Trp in proximity to Lys391.

### AmmB1 is a Leu-tRNA^Leu^ dependent PEARL enzyme

For the *bha* system, we were able to identify the next step after Gly addition by co-expression with the Gly oxidase BhaG. We therefore examined whether the same might be true for the *amm* system. Given that the glycine adduct **5** formed by AmmB_3_ is an isomer of **1**, and that the gene encoded by Amm14 (Figure 1A) is a putative glycine oxidase homolog of BhaG, we first co-expressed Amm14 with AmmA*-Trp, AmmC_1_, and AmmB_3_. However, we were not able to oxidize the glycine adduct in our co-expressions. Assuming that this might be a problem of expression of active Amm14, we investigated whether the glycine oxidase in the *bha* cluster (26% sequence identity to Amm14) could be used to catalyze 7-aminoquinone formation in the peptide. After including BhaG in the co-expression system described above, full conversion to the corresponding aminoquinone **6** was indeed observed (Figures S12 and 4B). High resolution MS/MS analysis showed that the modification was localized at the Trp. Thus, BhaG can oxidize Gly conjugated at either C5 or C7 of the oxidized indole of a C-terminal Trp.

We next investigated whether the formation of the aminoquinone in the *amm* pathway would unlock a new biosynthetic step en route towards the formation of the ammosamide core, similar to the assignment of BhaB_4_ as an Asn-tRNA ligase discussed above. Therefore, we co-expressed AmmA*-Trp with the Bha homologs of AmmC_1_ (BhaC_1_) and Amm14 (BhaG), AmmB_3_, and the two remaining PEARLs in the *amm* cluster, AmmB_1_ and AmmB_4_, in separate cultures. After purification of the full-length peptide and MALDI-TOF MS analysis, only co-expression with AmmB_1_ showed a distinct MALDI-TOF MS profile that showed the seemingly unreacted aminoquinone **6** as well as two additional peaks corresponding to an increase in mass of 42 and 85 Da compared to **6** (Figure 6A). High resolution MS/MS of the trypsin digested mixture revealed that the apparently unmodified aminoquinone corresponded to a product that had decreased in mass by 1 Da from the aminoquinone intermediate **6** (i.e. product **7**, Figure 6B) and the +42 and +85 Da products corresponded to acetylated **6** and bisacetylated reduced hydroquinone **7** as shown in Figure S13A. The most likely explanation is that product **7** is formed by addition of an amino acid by AmmB_1_ and subsequent conversion to the diaminoquinone (Figure 6B). In *E. coli* both starting peptide **6** and peptide **7** (or its initially formed hydroquinone derivative) are then acetylated by an unknown enzyme.

To first identify the cryptic amino acid adduct and avoid acetylation, an *in vitro* reaction was conducted that included AmmB_1_, a mixture of *E. coli* AARS and tRNAs as well as purified aminoquinone intermediate **6**. Using a mixture of all 20 L-amino acids labeled with ^15^N, we observed full incorporation of one ^15^N into the diaminoquinone **7** (Figure S14). We tested our initial hypothesis that considered the use of Gly-tRNA^Gly^ as the second amino group donor of the indole core. However, the corresponding *in vitro* reaction did not yield any product. Next, we performed *in vitro* reactions with the remaining amino acids split into three groups (Figure S14B) and found that the group containing leucine, isoleucine, valine, alanine, and proline gave the previously observed -1 Da product **7**. Individual amino acids from this group were then tested and only Leu reproduced the formation of the corresponding diaminoquinone core (Figure S14C). Thus, tRNA^Leu^ and N-terminally His_6_-tagged LeuRS from *E. coli* was expressed and purified. Using L-^15^N-leucine, LeuRS, and tRNA^Leu^ we observed the formation of the diaminoquinone product **7** with an increase in mass by 1 Da (Figure S14B). While the diaminoquinone product is observed in this reaction, we were not able to observe the anticipated leucine-adduct intermediate. This finding could indicate that AmmB_1_ has dual activity, first catalyzing the appendage of the leucine moiety to the Trp core and also the conversion to the observed product **7**. Two different pathways can be envisioned for the latter process that would produce either isovaleraldehyde or 4-methyl-2-oxovaleric acid from Leu (Figure S15A). To test this hypothesis, we incubated the *in vitro* reaction products with 2,4-dinitrophenylhydrazine. As shown in Figure S15A, we observed formation of the hydrazone adduct of 4-methyl-2-oxovaleric acid. This conclusion was confirmed by utilizing an authentic standard as well as several isotopically labeled L-leucine derivatives that all supported a hydrolytic mechanism, Figures S15B and S15C.

**Figure 6.**
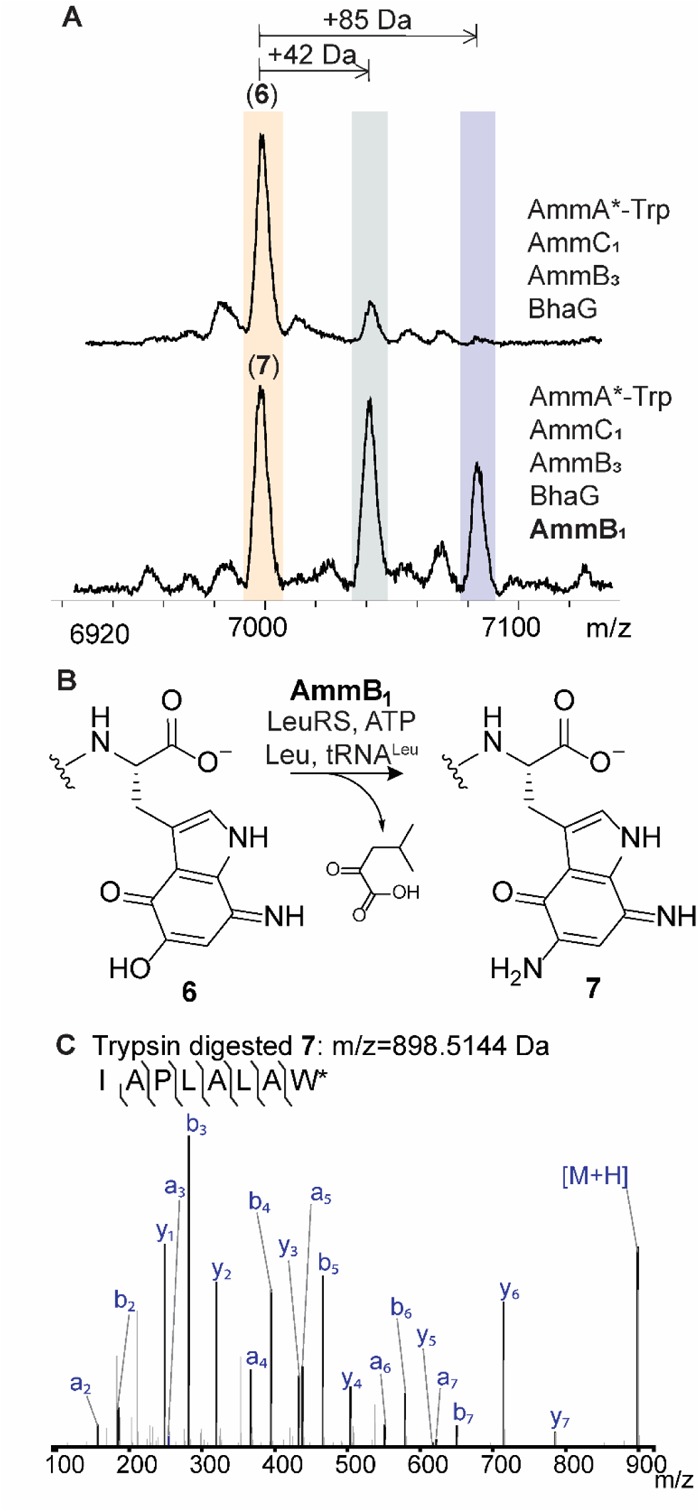
AmmB_1_ donates an amino group to the indole of intermediate **6**. (**A**) Inclusion of BhaB_4_ in the co-expression system that produces **6** results in the formation of three new products (Figure S13). (**B**) The reaction catalyzed by AmmB_1_. (**C**) ESI-MS/MS analysis of the C-terminal fragment upon trypsin digestion of peptide **7** (calculated m/z = 898.5144, observed m/z = 898.5181).

### The possible origin of the PEARL enzyme class

The discovery in 2015 that class I lanthipeptide dehydratases (LanBs) use glutamyl-tRNA to dehydrate Ser and Thr residues through a transesterification to form a glutamyl ester followed by glutamate elimination was surprising as no such biochemical transformations had been reported previously.^5^ Furthermore, the X-ray structure of a representative enzyme (NisB, UniProtKB P20103, PDB ID 4WD9) did not show any obvious structural homology with characterized enzymes to provide insights where this activity may have evolved from. A subset of LanB enzymes, called split LanBs, are encoded as two polypeptides, with one subunit catalyzing the glutamylation reaction and the second subunit catalyzing glutamate elimination.^9–11^ The discovery that the related PEARL enzymes that contain a glutamylation-like domain but not the glutamate elimination domain added amino acids to the C-termini of peptides in an amino acyl-tRNA and ATP dependent process was again surprising given that the lanthipeptide dehydratases do not utilize ATP. Here we used bioinformatic analysis as well as structure prediction by AlphaFold^12^ to try and answer some of these outstanding questions.

We constructed AlphaFold models of the PEARLs from the *bha* and *amm* clusters and used these models to gain insights into the putative catalytic region (Figure 7). The predicted PEARL structures revealed similar overall three-dimensional structures (e.g. Figure S16) with an apparent active site pocket resembling that of the crystallographically characterized split LanB, TbtB (UniProtKB D6Y502, PDB ID 6EC8) that catalyzes tRNA-dependent Ser glutamylation.^13^ In the TbtB structure, this pocket is occupied by a non-hydrolyzable substrate mimic of a 3’-glutamylated adenosine that interacts with several residues through H-bonding and π-π stacking interactions as shown in Figure S17. Two of the residues interacting with the adenine of the non-hydrolyzable mimic (Arg743 and Phe783 in TbtB) are also conserved in the PEARLs, as is a glutamic acid residue (Glu851, Figure 7). We hypothesize that in PEARLs this region may bind the adenosine of ATP and the 3’ terminus of a charged tRNA molecule, with the adenine moiety of both substrates interacting with the conserved Phe and Arg.

We compared the residues in this region of several AlphaFold-predicted PEARL structures as well as TglB (UniProtKB A0A8T8BZ30), a PEARL found in the thiaglutamate BGC that has been investigated by site-directed mutageneis.^2^ Two residues are structurally conserved in the predicted PEARL structures that are not present in TbtB, Asp542 and Arg803 (TglB numbering; Figure 7). Mutagenesis studies of these residues in TglB affected the phosphorylation step that activates the C-terminus of the precursor peptide required for amino acid condensation via aminoacyl-tRNA transfer (Figure 1A).^2^ Thus these prior mutagenesis studies on TglB are consistent with ATP binding in the same pocket as the non-hydrolyzable mimic of aminoacyl-tRNA.

**Figure 7.**
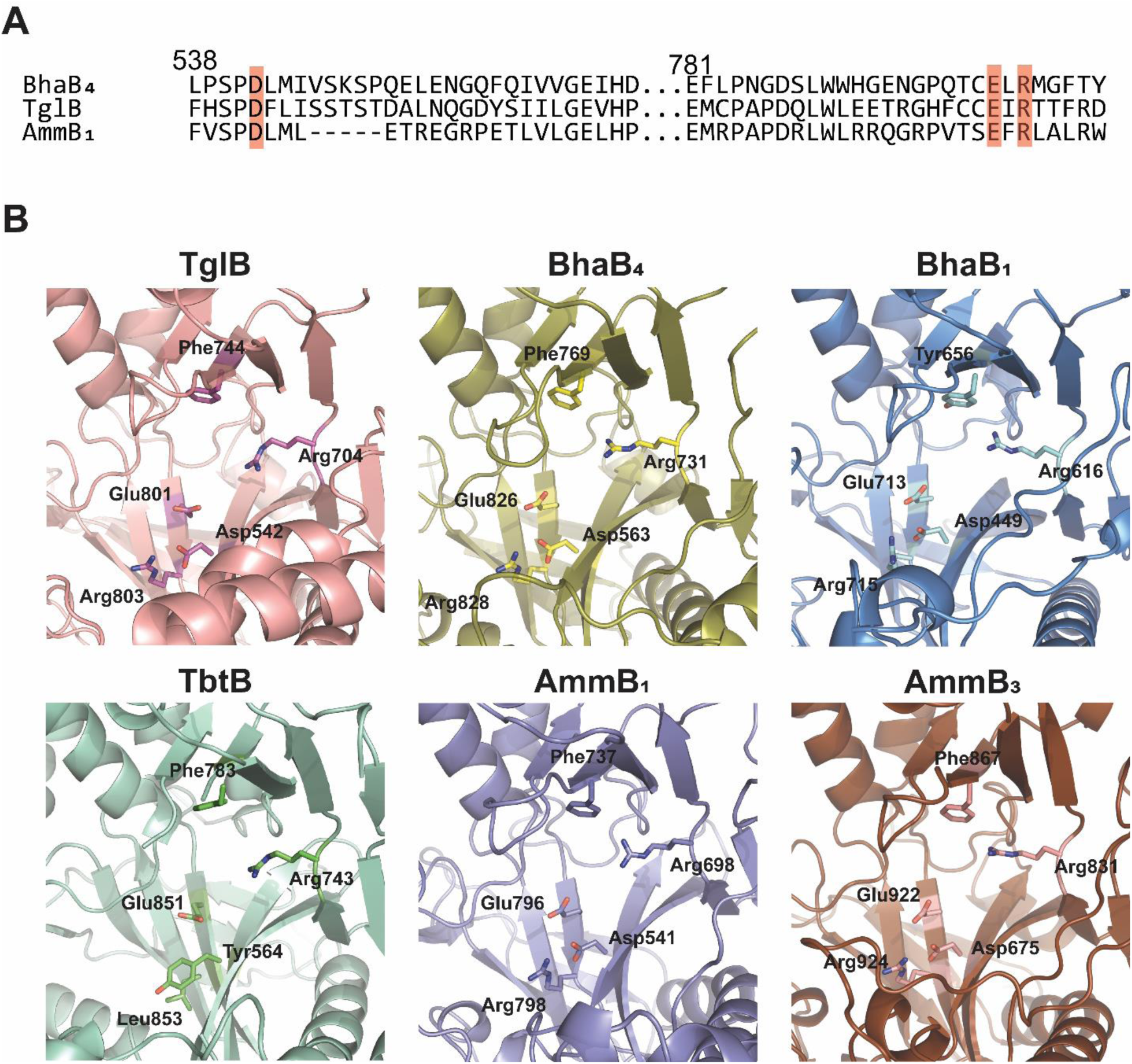
Conserved residues in PEARL enzymes versus TbtB, a member of the split LanB enzyme class. (A) Multiple Sequence Comparison by Log-Expectation (MUSCLE) alignment of three PEARL enzymes TglB, BhaB_4_, and AmmB_1_. Conserved residues (Asp542, Glu801, and Arg803; TglB numbering) located in the putative ATP-binding pocket are highlighted in red. (B) AlphaFold predicted structures of the PEARL TglB (pink) and split LanB TbtB (green) in comparison with two representative examples of the PEARL enzymes in the *bha* and *amm* clusters. The residues corresponding to Arg743, Phe783, and Glu851 in TbtB are conserved in all structures, but Asp542 and Arg803 of TglB are only conserved in the PEARLs and substituted by Tyr and Leu in split LanB enzymes.

Next, we searched for *bha* or *amm* sequence homologs using the Basic Local Alignment Search Tool (BLAST)^14^ with AmmB_4_ as query. Whereas the proteins with highest similarity scores were other PEARLs, split LanBs and full length LanBs, we also retrieved considerably shorter protein sequences (∼ half the size of PEARLs) with over 30% sequence identity (Figure S18). Intrigued by this result we submitted these sequences for AlphaFold analysis and the resulting structures to the Dali Server. The results of this 3D-based search indicated that the N-terminal portion of the shorter PEARL sequence resembled previously structurally characterized glutathionylspermidine synthase and glutathione synthetase, a founding member of the ATP-GRASP family^15^ (Figure S19). Like the PEARLs, these two enzymes catalyze condensation reactions at the C-terminus of a peptide in an ATP dependent manner involving first phosphorylation of the C-terminal carboxylate followed by attack of a nucleophilic amine onto the activated acylphosphate.

A comparison between glutathione (GSH) synthase and PEARL sequences and predicted structures revealed similar constellation of residues Glu281, Asp273, Arg210, and Asp208 in GSH synthase with the corresponding residues in the PEARLs (e.g. Glu564, Asp542, Arg803, and Glu801 in TglB, Figure S19). In the structure of GSH synthase bound to ADP (PDB 1GSA),^16,17^ Glu281 along with Asn283 mediates binding of two Mg^2+^ ions that coordinate to the β and α phosphate of ADP, and Asp273 binds to the Mg^2+^ that interacts with the α-phosphate. Arg210 and Asp208 are near the putative γ phosphate site (substituted by a sulfate group in the crystal structure) and the hydroxyl group in the ribose ring of ATP. In TglB, Glu564 was shown not to be essential for catalysis and Asn283 is replaced by a His residue that is highly conserved in PEARLs. These observations suggest that during the evolution of ATP-GRASP enzymes to PEARLs, some of the residues that interacted with ATP in the former were changed such that the latter could accommodate aminoacyl-tRNA in the same binding site. The split LanBs may have evolved from PEARLs by recruitment of the elimination domain that has evolutionary links to a number of other proteins (thiopeptide cyclases, LsrG epimerase),^18,19^ and the LanB lanthipeptide dehydratases arose from a subsequent gene fusion of the tRNA-dependent domain and the elimination domain.

## Discussion

After the previous unexpected discovery that PEARLs add glycine from Gly-tRNA to hydroxyquinones derived from Trp during the biosynthesis of pyrroloiminoquinone natural products,^4^ we proposed a chemically plausible route to ammosamide C that was efficient in terms of step-count (Figure S1), but did not use all of the biosynthetic genes in the BGC. In this study, we demonstrate the activity of some of these enzymes that allowed assignment of new enzymatic activities for two previously uncharacterized PEARLs in the *bha* and *amm* BGCs. Our data show that the amine groups in ammosamide do not all originate from glycine as we had anticipated, nor are they introduced using similar chemistry. Instead, for ammosamide we show that the nitrogen at C7 of the former indole is derived from glycine but the amine at C5 is derived from leucine. Furthermore, the mechanisms of cleavage of the N-Cα bonds of these two amino acids during the amino transfer process is quite different. For the introduction of the amine at position 7, a glycine oxidase is required to efficiently convert the initial glycine condensation product to the aminoquinone product, a reaction that produces glyoxylate. Conversely, for the introduction of the amino group at C5, a tautomerization and hydrolysis process appears to be operational (Figure S15).

The glycine oxidase enzyme is encoded in both the *amm* and *bha* BGCs, and its activity unlocked new PEARL reactions in both pathways. In the *amm* system, previous attempts to reconstitute the pathway beyond the glycine addition step catalyzed by AmmB_3_ stalled at intermediate **2’** (or its tautomer, Figure S10E), which we now consider a dead-end product. In the presence of the glycine oxidase, the aminoquinone intermediate was efficiently formed, which allowed the function of AmmB_1_ to be determined (Figure 4). Similarly, without the glycine oxidase, the *bha* pathway produced a mixture of products **1**-**3**, of which only the minor product (**3**) is a competent substrate for BhaB_4_. In addition to showing that the glycine oxidases are important for both pathways, our study revealed another important enzyme that assures the formation of the correct oxidation state of an intermediate for subsequent enzyme catalysis. The data show that the quinone reductase BhaH (and presumably its homolog Amm17 in the *amm* BGC), is required for BhaB_4_ activity by reducing the initially formed aminoquinone to the hydroquinone form. We suggest that the quinone reductases BhaH and Amm17 are involved in several steps of the pathway to reversibly interconvert quinone and hydroquinone forms of the former Trp indole. For instance, the initial BhaC_1_ and AmmC_1_ products are hydroxylated indoles (Figure 4), but for the PEARL chemistry catalyzed by AmmB_3_ and BhaB_5_ a vinylogous carboxylate is required for ATP-GRASP-like phosphorylation followed by an addition-elimination sequence with a tetrahedral intermediate. We suggest that quinone reductases may oxidize the BhaC_1_/AmmC_1_ product for use by the PEARL enzymes (Figure 4). Conversely, for the chlorinase Amm3 involved in ammosamide C biosynthesis,^20^ for which we and others have not yet been able to identify the exact substrate, a nucleophilic reduced oxidation state will be required.

In this study we identified the activities of two PEARLs. BhaB_4_ added Asn to the C-terminus of the peptide intermediate and AmmB_1_ added Leu to the oxidized aminoquinone core although the initial adduct was not detected and the addition is inferred from the isotope incorporation studies. Like all previously characterized PEARLs, these two activities were dependent on both ATP and the presence of the corresponding tRNA^Asn^ and tRNA^Leu^. Also similarly to previous studies, these enzymes worked with tRNA sequences from *E. coli* even though the enzymes originate from different bacterial phyla that use different tRNA sequences suggesting that it is mostly the amino acid that confers substrate specificity. Three PEARLs in the *bha* cluster remain to be assigned, as well as one PEARL in the *amm* cluster. Given the surprises that the biosynthetic pathways to pyrroloiminoquinones have provided thus far, we consider it perilous to propose a possible pathway from the currently accessed last intermediate to ammosamide C.

Ever since the mechanism of class I lanthipeptide dehydratases was shown to require glutamyl-tRNA, the evolutionary origin of these enzymes has remained enigmatic. Through the use of structure predicting tools like AlphaFold and bioinformatic analysis we have gained the first insights into both the three-dimensional features that may be important for PEARL catalysis and their potential evolutionary path. We suggest that PEARLs evolved from ATP-Grasp enzyme family members by recruiting a specialized nucleophile (aminoacyl-tRNA) to react with an activated acylphosphate substrate. It is possible that recruitment of this new substrate was facilitated by the similarities in structure between ATP and the conserved adenosine at the 3’ end of aminoacylated tRNA. Additional outstanding questions include the molecular basis for the amino acyl-tRNA specificities of PEARLs as well as the recognition of their peptide substrates.

## AUTHOR CONTRIBUTION

J.R.F. and W.A.V. designed research. J.R.F. performed experiments. J.R.F. and L.Z. performed NMR data acquisition and analysis. J.R.F and W.A.V. wrote the manuscript with input from all authors.

## AUTHOR INFORMATION

### Authors

Josseline Ramos Figueroa –Department of Chemistry and Howard Hughes Medical Institute, University of Illinois at Urbana-Champaign, Urbana, IL, United States.

Lingyang Zhu – School of Chemical Sciences NMR Laboratory, University of Illinois at Urbana-Champaign, Urbana, IL, United States.

### Funding

This work was supported in part by a grant from the National Institutes of Health (GM058822 to W.A.vdD.). W.A.vdD is an Investigator of the Howard Hughes Medical Institute. A Bruker UltrafleXtreme mass spectrometer used was purchased with support from the National Institutes of Health (S10 RR027109).

### Notes

The authors declare no competing financial interests.

## ACKNOWLEDGMENTS

HHMI lab heads have previously granted a nonexclusive CC BY 4.0 license to the public and a sublicensable license to HHMI in their research articles. Pursuant to those licenses, the author-accepted manuscript of this article can be made freely available under a CC BY 4.0 license immediately upon publication.

## Notes

### Competing Interest Statement

The authors have declared no competing interest.

## References

(1) Ting, C. P.; Funk, M. A.; Halaby, S. L.; Zhang, Z.; Gonen, T.; van der Donk, W. A. Use of a scaffold peptide in the biosynthesis of amino acid-derived natural products. Science 2019, 365, 280–4.

(2) Zhang, Z.; van der Donk, W. A. Nonribosomal peptide extension by a peptide amino-acyl tRNA ligase. J. Am. Chem. Soc. 2019, 141, 19625–33.

(3) Yu, Y.; van der Donk, W. A. Biosynthesis of 3-thia-α-amino acids on a carrier peptide. Proc. Natl. Acad. Sci. U. S. A. 2022, 119, e2205285119.

(4) Daniels, P. N.; Lee, H.; Splain, R. A.; Ting, C. P.; Zhu, L.; Zhao, X.; Moore, B. S.; van der Donk, W. A. A biosynthetic pathway to aromatic amines that uses glycyl-tRNA as nitrogen donor. Nat. Chem. 2022, 14, 71–7.

(5) Ortega, M. A.; Hao, Y.; Zhang, Q.; Walker, M. C.; van der Donk, W. A.; Nair, S. K. Structure and mechanism of the tRNA-dependent lantibiotic dehydratase NisB. Nature 2015, 517, 509–12.

(6) Daniels, P. N.; van der Donk, W. A. Substrate specificity of the flavoenzyme BhaC(1) that converts a C-Terminal Trp to a hydroxyquinone. Biochemistry 2023, 62, 378–87.

(7) Settembre, E. C.; Dorrestein, P. C.; Park, J.-H.; Augustine, A. M.; Begley, T. P.; Ealick, S. E. Structural and mechanistic studies on ThiO, a glycine oxidase essential for thiamin biosynthesis in *Bacillus subtilis*. Biochemistry 2003, 42, 2971–81.

(8) Evans, R.; O’Neill, M.; Pritzel, A.; Antropova, N.; Senior, A.; Green, T.; Žídek, A.; Bates, R.; Blackwell, S.; Yim, J., et al. Protein complex prediction with AlphaFold-Multimer. bioRxiv 2021, 2021.10.04.463034.

(9) Hudson, G. A.; Zhang, Z.; Tietz, J. I.; Mitchell, D. A.; van der Donk, W. A. In vitro biosynthesis of the core scaffold of the thiopeptide thiomuracin. J. Am. Chem. Soc. 2015, 137, 16012–5.

(10) Zhang, Z.; Hudson, G. A.; Mahanta, N.; Tietz, J. I.; van der Donk, W. A.; Mitchell, D. A. Biosynthetic timing and substrate specificity for the thiopeptide thiomuracin. J. Am. Chem. Soc. 2016, 138, 15511–4.

(11) Ozaki, T.; Kurokawa, Y.; Hayashi, S.; Oku, N.; Asamizu, S.; Igarashi, Y.; Onaka, H. Insights into the biosynthesis of dehydroalanines in goadsporin. ChemBioChem 2016, 17, 218–23.

(12) Jumper, J.; Evans, R.; Pritzel, A.; Green, T.; Figurnov, M.; Ronneberger, O.; Tunyasuvunakool, K.; Bates, R.; Žídek, A.; Potapenko, A., et al. Highly accurate protein structure prediction with AlphaFold. Nature 2021, 596, 583–9.

(13) Bothwell, I. R.; Cogan, D. P.; Kim, T.; Reinhardt, C. J.; van der Donk, W. A.; Nair, S. K. Characterization of glutamyl-tRNA-dependent dehydratases using nonreactive substrate mimics. Proc. Natl. Acad. Sci. U. S. A. 2019, 116, 17245–50.

(14) Altschul, S. F.; Gish, W.; Miller, W.; Myers, E. W.; Lipman, D. J. Basic local alignment search tool. J. Mol. Biol. 1990, 215, 403–10.

(15) Iyer, L. M.; Abhiman, S.; Maxwell Burroughs, A.; Aravind, L. Amidoligases with ATP-grasp, glutamine synthetase-like and acetyltransferase-like domains: synthesis of novel metabolites and peptide modifications of proteins. Mol. Biosyst. 2009, 5, 1636–60.

(16) Hara, T.; Kato, H.; Katsube, Y.; Oda, J. i. A pseudo-Michaelis quaternary complex in the reverse reaction of a ligase: structure of *Escherichia coli* b glutathione synthetase complexed with adp, glutathione, and sulfate at 2.0 Å resolution. Biochemistry 1996, 35, 11967–74.

(17) Galperin, M. Y.; Koonin, E. V. A diverse superfamily of enzymes with ATP-dependent carboxylate-amine/thiol ligase activity. Protein Sci. 1997, 6, 2639–43.

18. Cogan, D. P.; Hudson, G. A.; Zhang, Z.; Pogorelov, T. V.; van der Donk, W. A.; Mitchell, D. A.; Nair, S. K. Structural insights into enzymatic [4+2] aza-cycloaddition in thiopeptide antibiotic biosynthesis. Proc. Natl. Acad. Sci. U. S. A. 2017, 114, 12928–33.

(19) Marques, J. C.; Lamosa, P.; Russell, C.; Ventura, R.; Maycock, C.; Semmelhack, M. F.; Miller, S. T.; Xavier, K. B. Processing the interspecies quorum-sensing signal autoinducer-2 (AI-2): characterization of phospho-(S)-4,5-dihydroxy-2,3-pentanedione isomerization by LsrG protein. J. Biol. Chem. 2011, 286, 18331–43.

(20) Jordan, P. A.; Moore, B. S. Biosynthetic pathway connects cryptic ribosomally synthesized posttranslationally modified peptide genes with pyrroloquinoline alkaloids. Cell Chem. Biol. 2016, 23, 1504–14.

