## Supporting Information Figures for "Unexpected transformations during pyrroloiminoquinone biosynthesis"

### General materials and methods

Molecular biology experiments were performed using reagents from New England Biolabs, Thermo Fisher Scientific, or Sigma-Aldrich. Plasmid maintenance and protein overexpression were conducted using *Escherichia coli* DH10β and BL21 (DE3) strains. pRSFDuet-1 and pCDFDuet-1 were obtained from Novagen. MALDI-TOF-MS data acquisition was performed using a Bruker UltrafleXtreme mass spectrometer (Bruker Daltonics) in reflector positive mode at the University of Illinois School of Chemical Sciences Mass Spectrometry Laboratory. A commercial mixture of 9:1 2,5-dihydroxybenzoic acid (DHB) and 2-hydroxy-5-methoxybenzoic acid (Super-DHB or SDHB) solution with a stock concentration of 25 mg/mL was routinely mixed with samples in a 1 to 1 ratio and air-dried before MALDI-TOF-MS analysis. MS1 and MS2 data acquisition was performed using an Agilent qTOF instrument equipped with an UPLC system. 2D NMR experiments were recorded using an Agilent 600 MHz spectrometer in 100% $d_6$-DMSO. Ni-NTA resin for peptide and protein purifications was obtained from Thermofisher.

### Molecular Biology Techniques

Plasmid DNA was prepared using a MiniPrep kit (Qiagen) following the manufacturer's instructions from *E. coli* DH10β cells. Plasmids were manually designed and prepared by Gibson assembly using NEBuilder HiFi DNA Assembly Master Mix. The vector primer design usually incorporated overhangs complementary to the gene of interest (Table S1). PCR amplification was performed using Q5 polymerase, dNTPs and Q5 reaction buffer from New England Biolabs. Designed primers for cloning and in vitro transcription were acquired from Integrated DNA Technologies (Table S1). Genes encoding *E. coli* asparagine $tRNA^{Asn}$ and leucine $tRNA^{Leu}$ ligase were obtained from TwistBioscience (Table S1).

### Peptide expression and purification

Peptide expression was carried out by growing co-transformed *E. coli* BL21 (DE3) cells expressing N-terminal $His_6$-tagged BhaA_AW and untagged BhaB5 from pACYCDuet-1(MCSI:His6-BhaA_AW and MCSII:BhaB5), untagged BhaC1 from pCDFDuet-1 (MCSI:BhaC1) and additional enzymes BhaG, BhaB2, BhaB3, BhaB4, BhaB6, BhaF, BhaC2, BhaC3, BhaC4, and BhaH from the corresponding pET28 plasmids. Cells were grown with 25 µg/mL chloramphenicol, 100 µg/mL spectinomycin, and 50 µg/mL kanamycin in Luria Bertani Broth. Cultures were shaken at 190 RPM at 37 $^{o}$C for 3 h following inoculation with IPTG to a final concentration of 0.4 mM when OD600 reached 0.6-0.8. After incubation for 18 h overnight at 18 $^{o}$C, cells were harvested by centrifugation at 5,200 ×*g*. Cell pellets were stored in a -80 $^{o}$C freezer for future use or resuspended in denaturing buffer composed of 6 M guanidinium hydrochloride, 50 mM 4-(2-hydroxyethyl)-1-piperazineethanesulfonic acid (HEPES), pH 7.6. Next, resuspended cells were disrupted by sonication using repeated cycles of 2 s on and 5 s off. After centrifugation at 50,000 ×*g*, clarified lysate was passed through Ni-NTA resin to retain the His-tagged peptide. The column was washed with 20 column volumes (CV) of wash buffer composed of 25 mM imidazole, 300 mM NaCl, and 50 mM HEPES at pH 7.6. The His-tagged peptide was then eluted with 5 CV of elution buffer composed of 200 mM imidazole, 300 mM NaCl, and 50 mM HEPES at pH 7.6. The peptide was then purified by HPLC or desalted using a C18 ziptip and eluted into 1.5 µL of 25 mg/mL of a solution of SDHB dissolved in 80% $CH_3CN$ (ACN)/20% $H_2O$/0.1% HCOOH (FA) for MALDI-TOF-MS analysis.

**High-performance liquid chromatography (HPLC) purification of isolated peptide**

Peptides obtained from Ni-resin purification were acidified with a 10% trifluoroacetic acid (TFA) solution to a final concentration of 0.1 to 2% TFA to precipitate any coeluting protein from the overexpression protocol. The sample was then passed through a 0.45 µm syringe filter and injected directly into an Agilent 1260 Infinity II HPLC instrument for purification. The peptide was purified using a preparative scale column, VP NUCLEODUR C18 HTec, 5 µm, 250x10 mm with the following gradient: 0-5 min isocratic 2% ACN, 5-40 min 2%-60% ACN, 40-42 min 60%-100% ACN; mobile phase buffered with 0.1% TFA. Fractions were collected manually and analyzed by MALDI-TOF MS by combining 1 to 1 with a 25 mg/mL solution of SDHB. Fractions containing the desired peptide were combined and lyophilized. The dried solids were then used for in vitro assays, further purification on an analytical scale column, or proteolytic digestion for LC-MS or NMR analysis.

When an additional round of purification was required, peptides were redissolved in water, passed through a 0.45 µm filter, and injected directly into an analytical scale Agilent 1260 Infinity HPLC. A Vydac 218TP C18 column, 5 µm, 250x4.6 mm was used for peptide purification with the following gradient: 0-10 min isocratic 2% ACN, 10-30 min 2%-60% ACN, 30-32 min 60%-100% ACN; mobile phases contained 0.1% TFA. The desired peptide was identified by MALDI-TOF MS analysis performed on each collected fraction.

When the samples were used for NMR experiments or LC-MS/MS analysis, the HPLC purified peptides were subjected to trypsin proteolytic cleavage, acidified to 0.1 to 1% TFA, filtered through a 0.45 µm membrane or centrifuged at 20,000 ×*g* for 5 min, and purified by HPLC. The tryptic fragment containing the C-terminal modifications was then eluted with the same column and gradient as indicated above for analytical scale purification. Fractions were collected manually and submitted for MALDI-TOF MS analysis for peptide fragment identification.

**BhaG expression and purification**

Protein expression was carried out by growing *E. coli* BL21 (DE3) cells expressing the N-terminal $His_6$-tagged BhaG from pET28-BhaG. Cells were grown with 50 µg/mL kanamycin in Luria Bertani Broth. Cultures were shaken at 190 RPM at 37 °C for 3 h following inoculation with IPTG to a final concentration of 0.4 mM when OD600 reached 0.6-0.8 units. After incubation for 18 h overnight at 18 °C, cells were harvested by centrifugation at 5,200 ×*g*. Cell pellets were stored in a -80 °C freezer for future use or resuspended in non-denaturing buffer composed of 20 mM $NaH_2PO_4$, 500 mM NaCl, and 50 mM imidazole at pH 7.5 with 5% glycerol. Next, resuspended cells were disrupted by sonication with repeated cycles of 2 s on and 5 s off. After centrifugation at 50,000 ×*g*, the clarified lysate was passed through Ni-NTA resin, previously equilibrated with the same initial buffer, to retain the His-tagged protein. The column was washed further with 20 CV of the initial buffer. Then, the His-tagged protein was eluted with 5 CV of elution buffer composed of 20 mM $NaH_2PO_4$, 500 mM NaCl, and 200 mM imidazole at pH 7.5 with 5% glycerol. After that, a previously equilibrated PD-10 column was used to buffer exchange the protein solution to 50 mM $K_3PO_4$ and 5% glycerol at pH 8.0. The flowthrough was further concentrated using an Amicon centrifugal device with a 30 kDa MWCO. A slight precipitation of the yellow protein was observed and which was redissolved by adding glycerol in a 1 to 1 ratio.

**$BhaB_4$ and $AmmB_1$ expression and purification**

Protein expression was carried out by growing transformed *E. coli* BL21 (DE3) cells expressing the N-terminal $His_6$-tagged PEARL $BhaB_4$ or $AmmB_1$ from the corresponding pET28-$BhaB_4$ or pET28-$AmmB_1$ plasmids. Cells were grown with 50 µg/mL kanamycin in Luria Bertani Broth. Cultures were shaken at 190 RPM at 37 °C for 3 h following inoculation with IPTG to a final concentration of 0.4 mM when OD600 reached 0.6-0.8 units. After incubation for 18 h overnight at 18 °C, cells were harvested by centrifugation at 5,200 ×*g*. Pellets were stored in a -80 °C freezer for future use or resuspended in non-denaturing buffer composed of 20 mM HEPES, 300 mM NaCl, and 50 mM imidazole at pH 7.5 with 5% glycerol. Next, resuspended cells were disrupted by sonication with repeated cycles of 2 s on and 5 s off. After centrifugation at 50,000 ×*g*, clarified lysate was passed through Ni-NTA resin, previously equilibrated with the same initial buffer, to retain the His-tagged protein. The column was washed further with 20 CV of the initial buffer. Then, the His-tagged protein was eluted with 5 CV of elution buffer composed of 20 mM HEPES, 500 mM NaCl, and 200 mM imidazole at pH 7.5 with 5% glycerol. After that, an Amicon with a 50 kDa cutoff was used to buffer exchange the protein solution to 50 mM HEPES and 5% glycerol at pH 7.5 and further concentrated to up to 30 mg/mL protein solution. Aliquots were taken and flash-frozen in liquid nitrogen before storing the protein solutions at -80 °C.

### In vitro Transcription of *E. coli* $tRNA^{Asn}$ and $tRNA^{Leu}$

tRNA was obtained as described previously with minor modifications.[1] First, double-stranded DNA (dsDNA) used as a template for transcription was produced using DNA Polymerase I, Large (Klenow) fragment according to the manufacturer's protocol. Briefly, forward, and reverse primers encompassing the desired nucleotide sequence and with a ten-nucleotide overlap (Table S1) were dissolved to a stock solution of 100 µM for each primer. Primers, dNTPs, and NEBuffer™ 2 (10X) were combined to a final concentration of 4 µM, 33 µM, and 1X, respectively. Then, 3 U of Klenow fragment was added while keeping all components on ice. After that, the mixture was incubated at 25 °C for 15 min. The reaction was stopped by adding EDTA to 10 mM final concentration and incubated at 75 °C for 20 min. The reaction was usually set up in four individual 200-µL tubes. After quenching, all reactions were pull together and the DNA was precipitated by adding 2.5 volumes of ice-cold 100% ethanol. The mixture was left at -80 °C for 1 h and then centrifuged at 20,000 ×*g* for 30 min at 4 °C. The supernatant was discarded and the solid was washed with 75% EtOH twice. The resulting pellet was then air dried for 15 min and re-dissolved in nuclease free water (~20 µL). After concentration quantification by nanodrop, the solution was aliquoted and stored at -20 °C for future use. In vitro transcription was performed using HiScribe® T7 High Yield RNA Synthesis Kit according to the manufacturer's protocol for short transcripts (< 0.3 kb). Briefly, nuclease-free water and kit components were mixed to a final concentration of 1X Reaction Buffer, 7.5 mM ATP, 7.5 mM GTP, 7.5 mM CTP, 7.5 mM UTP, and 1 µg of the dsDNA template in a final volume of 20 µL. Then 1.5 µL of T7 RNA Polymerase Mix was added and the mixture was incubated for 16 h at

37 °C. Four reactions were set up at the same time and pooled together for use in the next step. DNA template was removed by adding 8 µL of DNase I (RNase-free) using 1X DNase I Buffer and 280 µL of nuclease-free water. The mixture was incubated for 15 min at 37 °C. Then, the mixture was extracted with an equal volume of acidic phenol:$CHCl_3$ using a vortex-mixer for 1 min and centrifuged at 20,000 ×*g* for 5 min at 4 °C. The top phase was then carefully removed and mixed with an equal volume of a mixture of $CHCl_3$ and isoamyl alcohol (24:1), vortexed, and centrifuged again for 5 min to separate the phases at 4 °C. The top phase was removed and mixed with 2.5 volumes of cold 100 % EtOH to precipitate tRNA on ice. The solid was then separated by centrifugation at 20,000 ×*g* for 15 min at 4 °C. The supernatant was discarded, and the pellet washed twice with ice-cold 75% EtOH. The tRNA was air dried and resuspended in approximately 20 µL of 2 mM NaOAc pH 5.2. The concentration was determined by nanodrop and adjusted to 1 to 5 µg/µL before aliquoting and storing at -20 °C for future use.

**In vitro assay of BhaG and phenylhydrazine derivatization**

A 50-µL solution containing a final concentration of 25 mM HEPES pH 7.6, 1 mM TCEP pH 7.6, 5 µM BhaG, 0.5 mM $MgCl_2$, and 30 µM of His-tagged Intermediate **1** was incubated for 2 h at 37 °C. After that time, the reaction mixture was combined with 50 µL of Ni-resin slurry, vortexed, and centrifuged at 20,000 xg for 10 min. The supernatant (~80 µL) was removed and combined with 50 µL of a solution of 20 mM freshly prepared phenylhydrazine solution. The resulting solution was incubated for 2 h at 37 °C and then submitted for LC-MS analysis. The resin was then treated with 50 µL of 200 mM imidazole solution to elute the bound modified product of the BhaG reaction (intermediate 3) and centrifuged at 20,000 xg for 10 min. The resulting supernatant was transferred to an Amicon 3 kDa MW cutoff to buffer exchange the high concentration of imidazole to 50 mM HEPES pH 7.6 buffer. The resulting solution was treated with 2 µg of trypsin protease and incubated for 2 h at 37 °C. Finally, the digested peptide solution was desalted using a Pierce™ C18 column and the sample submitted for ESI-MS/MS analysis.

**In vitro assay of $BhaB_4$ and $AmmB_1$**

In a 200 µL tube, a solution containing a final concentration of 50 mM HEPES, 25 mM KCl, and 15 mM $MgCl_2$ pH 7.6, 2 mM DTT, 5 mM amino acid (asparagine or leucine for $BhaB_4$ or $AmmB_1$, respectively, or a mix of $^{15}$N-labeled L-amino acids), 6 mM ATP, 10 U Thermostable Inorganic

Pyrophosphatase (TIPP), 10 U Superase RNAse inhibitor, 60-200 ng/µL of tRNA (*E. coli* mix, $tRNA^{Asn}$, or $tRNA^{Leu}$), 20 µM of the corresponding aminoacyl-tRNA synthetase AsnRS or LeuRS, 5 µM of PEARL $BhaB_4$ or $AmmB_1$, and 15 µM of peptide substrate was incubated for 4 h at 37 °C in a heating block. After this time, the modified His-tagged peptide was isolated using Ni agarose resin. The eluted His-tagged peptide was buffer exchanged into 50 mM HEPES pH 7.5 and treated with trypsin or chymotrypsin for ESI-MS/MS analysis.

For the reaction with $AmmB_1$, 2,4-dinitrophenylhydrazine (2,4-DNP) was used to determine whether isovaleraldehyde or 4-methyl-2-oxopentanoic acid was formed after PEARL-catalyzed leucine addition to intermediate **6**. After the 4 h incubation with $AmmB_1$ as indicated above, the reaction mixture was diluted with an equal amount of water and 2,4-DNP was added to a final concentration of 2 mM. The resulting mixture was incubated at 37 °C for 16 h and then centrifuged through an Amicon 3 kDa MW cutoff for 10 min. The flowthrough was used for ESI-MS/MS analysis.

#### LC-MS analysis of the products of reactions with BhaG, $BhaB_4$, or $AmmB_1$

After trypsin digestion, the solutions containing peptide fragments were desalted using a Pierce™ C18 spin column. Briefly, the spin column was activated with 200 µL of 60% ACN with 0.1% formic acid (FA) and rinsed three times with 200 µL of 0.1% FA. The sample was passed through the equilibrated column and washed three times with 200 µL of 0.1% FA. Then, the peptide fragments were eluted using 25 µL of 60% ACN with 0.1% FA, twice. The eluant was lyophilized, redissolved in 25-50 µL of water, and passed through a 0.45 µm membrane to remove any insoluble material. The sample was then injected to an Agilent 1260 Infinity II mass spectrometer with a 6545 LC/Q-TOF detector. A kinetex® 2.6 µm C8 100 Å column, 150x2.1 mm, was used for peptide separation using the following gradient: 0-3 min isocratic 5% ACN, 3-13 min 5%-95% ACN, 13-15 min 95%-5% ACN; mobile phases contained 0.1% FA. A DualAJS ESI detector was utilized with the following parameters, Gas Temp of 325 °C, Drying Gas of 13 L/min, Nebulizer of 35 psi, Sheath Gas Temp of 275 °C, Sheath Gas Flow 12 L/min, Nozzle Voltage of 500 V and MS-TOF Fragmentor 175 V.

#### NMR characterization of the peptides modified by $BhaB_4$ and $AmmB_3$

Trypsin generated fragments of the products of $BhaB_4$, $AmmB_3$, and $AmmB_1$ were dissolved in DMSO-$d_6$ and transferred into a 5 mm Shigemi tube DMSO-$d_6$ matched solvent and bottom length of 8 mm compatible with Bruker. The sample was then submitted for NMR analysis on a 600 MHz Bruker spectrometer equipped with a Prodigy probe and a SampleXpress autosampler.

**Table S1. DNA sequences for primers and enzymes used in this study**

| | |
|---|---|
| tRNA_Ec_Leu.R | mUmGGTGCTGATAGGCAGATTCGAACTGCCGaCCTCACCCTTACCAAGGGTGCGCTCTACCA |
| tRNA_Ec_Leu.F | AATTCCTGCAGTAATACGACTCACTATAGCTGATATAGCTCAGTTGGTAGAGCGCACCCT |
| pRSF_for_LeuS.F | TTACGTACCAGGTAAACTCCTCAATCTGGTCGTTGGCTAATGCAGGTCGACAAGCTTGCG |
| pRSF_for_LeuS.R | GGATTCTATCTCTTCCGGGCGGTATTGCTCTTGCATCGGATCCTGGCTGTGGTGATGATG |
| pCDF-BhaC1_for_BhaG.R | GTGAAAGGACGGAGAGCTCGTAGCATATGTATATCTCCTTCTTATACTTAACTAATATAC |
| pCDF-BhaC1_for_BhaG.F | CGTCTAGTTTTGTTCGTTCGGATAACATGATATCGGCCGGCCAC |
| BhaG_for_pCDF-BhaC1.R | CTTGTCGACCTGCAGGCGCGCCGAGCTCGAATTCctacctcctttcaaccatcgcaaagc |
| BhaG_for_pCDF-BhaC1.F | TAACTTTAATAAGGAGATATACCATGGatgaaagagagatttgacctatgcataataggt |
| pCDF_for_BhaC1.R | GTGAAAGGACGGAGAGCTCGTAGCATATGTATATCTCCTTCTTATACTTAACTAATATAC |
| pCDF_for_BhaC1.F | CGTCTAGTTTTGTTCGTTCGGATAACATGATATCGGCCGGCCAC |
| BhaC1_for_pCDFmcs1.R | CGACGTCAGCGATCGCGTGGCCGGCCGATATCATGTTATCCGAACGAACAAAACTAGACG |
| BhaC1_for_pCDFmcs1.F | GTATATTAGTTAAGTATAAGAAGGAGATATACATATGCTACGAGCTCTCCGTCCTTTCAC |
| pRSF_for_AsnS.F | CGTACACCTCGCAATGCATCCTTCTAAATAATGCTTAAGTCGAACAGAAAGTAATCG |
| pRSF_for_AsnS.R | CACGGCCTTGCAGAACATCTGCAACCGGGACAACGCTCATCGGATCCTGGCTGTGGTG |
| tRNA_Ec_Asn.F | AATTCCTGCAGTAATACGACTCACTATATCCTCTGTAGTTCAGTCGGTAGAACGGCGGA |
| tRNA_Ec_Asn.R | mUmGGCTCCTCTGACTGGACTCGAACCAGTGaCATACGGATTAACAGTCCGCCGTTCTACCG |
| *E. coli* Leucyl-tRNA Synthetase (LeuRS) | ATGCAAGAGCAATACCGCCCGGAAGAGATAGAATCCAAAGTACAGCTTCATTGGGATGAGAAGCGCACATTTGAAGTAACCGAAGACGAGAGCAAAGAGAAGTATTACTGCCTGTCTATGCTTCCCTATCCTTCTGGTCGACTACACATGGGCCACGTACGTAACTACACCATCGGTGACGTGATCGCCCGCTACCAGCGTATGCTGGGCAAAAACGTCCTGCAGCCGATCGGCTGGGACGCGTTTGGTCTGCCTGCGGAAGGCGCGGCGGTGAAAAACAACACCGCTCCGGCACCGTGGACGTACGACAACATCGCGTATATGAAAAACCAGCTCAAAATGCTGGGCTTTGGTTATGACTGGAGCCGCGAGCTGGCAACCTGTACGCCGGAATACTACCGTTGGGAACAGAAATTCTTCACCGAGCTGTATAAAAAAGGCCTGGTATATAAGAAGACTTCTGCGGTCAACTGGTGCCCGAACGACCAG |

| | |
|---|---|
| | ACCGTACTGGCGAACGAACAAGTTATCGACGGCTGCTGCTGGCGCTGC<br>GATACCAAAGTTGAACGTAAAGAGATCCCGCAGTGGTTTATCAAAATCAC<br>TGCTTACGCTGACGAGCTGCTCAACGATCTGGATAAACTGGATCACTGG<br>CCAGACACCGTTAAAACCATGCAGCGTAACTGGATCGGTCGTTCCGAAG<br>GCGTGGAGATCACCTTCAACGTTAACGACTATGACAACACGCTGACCGT<br>TTACACTACCCGCCCGGACACCTTTATGGGTTGTACCTACCTGGCGGTA<br>GCTGCGGGTCATCCGCTGGCGCAGAAAGCGGCGGAAAATAATCCTGAA<br>CTGGCGGCCTTTATTGACGAATGCCGTAACACCAAAGTTGCCGAAGCTG<br>AAATGGCGACGATGGAGAAAAAAGGCGTCGATACTGGCTTTAAAGCGGT<br>TCACCCATTAACGGGCGAAGAAATTCCCGTTTGGGCAGCAAACTTCGTA<br>TTGATGGAGTACGGCACGGGCGCAGTTATGGCGGTACCGGGGCACGAC<br>CAGCGCGACTACGAGTTTGCCTCTAAATACGGCCTGAACATCAAACCGG<br>TTATCCTGGCAGCTGACGGCTCTGAGCCAGATCTTTCTCAGCAAGCCCT<br>GACTGAAAAAGGCGTGCTGTTCAACTCTGGCGAGTTCAACGGTCTTGAC<br>CATGAAGCGGCCTTCAACGCCATCGCCGATAAACTGACTGCGATGGGC<br>GTTGGCGAGCGTAAAGTGAACTACCGCCTGCGCGACTGGGGTGTTTCC<br>CGTCAGCGTTACTGGGGCGCGCCGATTCCGATGGTGACGCTGGAAGAC<br>GGTACCGTAATGCCGACCCCGGACGACCAGCTGCCGGTGATCCTGCCG<br>GAAGATGTGGTAATGGACGGCATTACCAGCCCGATTAAAGCAGATCCGG<br>AGTGGGCGAAAACTACCGTTAACGGTATGCCAGCACTGCGTGAAACCGA<br>CACTTTCGACACCTTTATGGAGTCCTCCTGGTACTATGCGCGCTACACTT<br>GCCCGCAGTACAAAGAAGGTATGCTGGATTCCGAAGCGGCTAACTACTG<br>GCTGCCGGTGGATATCTACATTGGTGGTATTGAACACGCCATTATGCAC<br>CTGCTCTACTTCCGCTTCTTCCACAAACTGATGCGTGATGCAGGCATGG<br>TGAACTCTGACGAACCAGCGAAACAGTTGCTGTGTCAGGGTATGGTGCT<br>GGCAGATGCCTTCTACTATGTTGGCGAAAACGGCGAACGTAACTGGGTT<br>TCCCCGGTTGATGCTATCGTTGAACGTGACGAGAAAGGCCGTATCGTGA<br>AAGCGAAAGATGCGGCAGGCCATGAACTGGTTTATACCGGCATGAGCAA<br>AATGTCCAAGTCGAAGAACAACGGTATCGACCCGCAGGTGATGGTTGAA<br>CGTTACGGCGCGGACACCGTTCGTCTGTTTATGATGTTTGCTTCTCCGG<br>CTGATATGACTCTCGAATGGCAGGAATCCGGTGTGGAAGGGGCTAACC<br>GCTTCCTGAAACGTGTCTGGAAACTGGTTTACGAGCACACAGCAAAAGG<br>TGATGTTGCGGCACTGAACGTTGATGCGCTGACTGAAAATCAGAAAGCG<br>CTGCGTCGCGATGTGCATAAAACGATCGCTAAAGTGACCGATGATATCG<br>GCCGTCGTCAGACCTTCAACACCGCAATTGCGGCGATTATGGAGCTGAT<br>GAACAAACTGGCGAAAGCACCAACCGATGGCGAGCAGGATCGCGCTCT<br>GATGCAGGAAGCACTGCTGGCCGTTGTCCGTATGCTTAACCCGTTCACC<br>CCGCACATCTGCTTCACGCTGTGGCAGGAACTGAAAGGCGAAGGCGAT<br>ATCGACAACGCGCCGTGGCCGGTTGCTGACGAAAAAGCGATGGTGGAA<br>GACTCCACGCTGGTCGTGGTGCAGGTTAACGGTAAAGTCCGTGCCAAAA<br>TCACCGTTCCGGTGGACGCAACGGAAGAACAGGTTCGCGAACGTGCTG<br>GCCAGGAACATCTGGTAGCAAAATATCTTGATGGCGTTACTGTACGTAAA<br>GTGATTTACGTACCAGGTAAACTCCTCAATCTGGTCGTTGGCTAA |
| *E. coli* Asparaginyl-tRNA Synthetase (AsnRS) | ATGAGCGTTGTCCCGGTTGCAGATGTTCTGCAAGGCCGTGTTGCTGTCG<br>ATTCGGAGGTTACCGTGCGTGGGTGGGTGCGTACCAGACGTGATAGTA<br>AAGCAGGTATTAGTTTCCTTGCGGTATATGATGGGTCGTGTTTTGATCCT<br>GTTCAAGCAGTGATTAATAATTCACTGCCAAATTACAATGAGGATGTACT<br>GCGTCTTACAACAGGTTGTTCTGTGATTGTAACTGGCAAGGTTGTCGCG<br>AGTCCTGGTCAAGGTCAGCAATTCGAGATTCAGGCGTCTAAAGTGGAAG<br>TTGCAGGGTGGGTGGAAGATCCGGACACCTATCCAATGGCAGCCAAAC<br>GGCACAGTATTGAATACCTGCGCGAGGTGGCCCACCTCCGTCCCCGCA |

| | |
|---|---|
| | CAAATCTGATCGGCGCTGTCGCTCGCGTACGCCACACCCTGGCTCAGG<br>CACTGCACCGTTTTTTCAATGAACAGGGATTTTTTTGGGTAAGCACACCG<br>CTGATTACCGCGTCGGATACCGAAGGGGCTGGCGAGATGTTTCGAGTTT<br>CAACACTGGATTTAGAAAATCTGCCACGCAACGATCAGGGCAAAGTCGA<br>TTTTGATAAAGACTTCTTTGGAAAAGAGAGTTTTTTAACCGTATCGGGGC<br>AGCTGAATGGAGAAACGTATGCATGCGCCTTGTCTAAAATCTACACATTT<br>GGACCCACGTTTCGTGCGGAGAACTCAAATACGTCTCGACACCTCGCAG<br>AATTTTGGATGCTTGAACCGGAGGTAGCGTTCGCCAACCTCAACGATAT<br>AGCAGGCCTGGCAGAAGCAATGCTCAAATATGTTTTTAAAGCGGTACTG<br>GAAGAGCGCGCTGATGACATGAAATTCTTTGCTGAACGGGTAGATAAAG<br>ATGCTGTGTCACGGCTGGAACGCTTCATTGAGGCAGATTTTGCCCAGGT<br>CGACTACACGGACGCCGTCACAATCCTGGAAAATTGTGGCCGAAAATTT<br>GAAAACCCCGTCTACTGGGGTGTTGATCTGTCTTCTGAACACGAAAGAT<br>ATCTTGCAGAAGAGCATTTCAAAGCCCCGGTAGTGGTAAAGAATTATCCA<br>AAAGATATTAAAGCGTTTTACATGCGCCTGAACGAAGATGGGAAGACGG<br>TAGCTGCGATGGACGTCTTGGCACCGGGTATTGGAGAAATCATTGGGG<br>GTTCGCAACGTGAGGAACGGTTGGATGTGCTCGATGAACGCATGTTAGA<br>AATGGGGTTAAATAAAGAGGATTATTGGTGGTATCGAGACCTTCGGCGT<br>TACGGGACTGTGCCGCATAGTGGCTTCGGTTTAGGTTTCGAACGGCTGA<br>TTGCTTATGTTACGGGGGTCCAAAATGTGCGAGACGTTATCCCTTTCCCA<br>CGTACACCTCGCAATGCATCCTTCTAA |

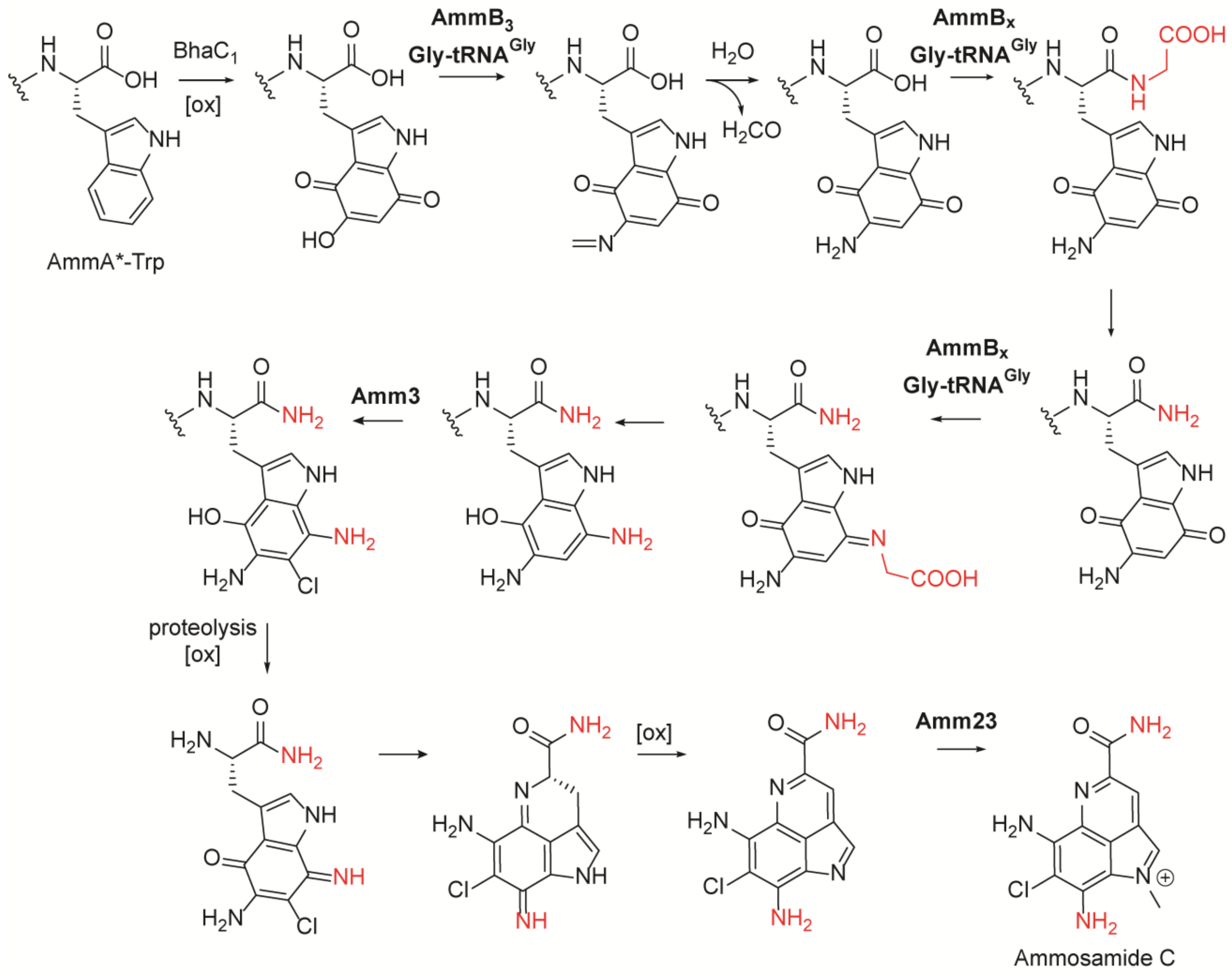


**Figure S1.** Biosynthetic pathway towards ammosamide C proposed in a previous study[2] postulating that a PEARL in the *amm* cluster would install the second amino group (red font) through a glycine addition to the oxidized C-terminal Trp. Similarly, the amide group nitrogen in the final product (also red font) was proposed to be derived from Gly-tRNA. The order of the various proposed steps could be different than depicted.

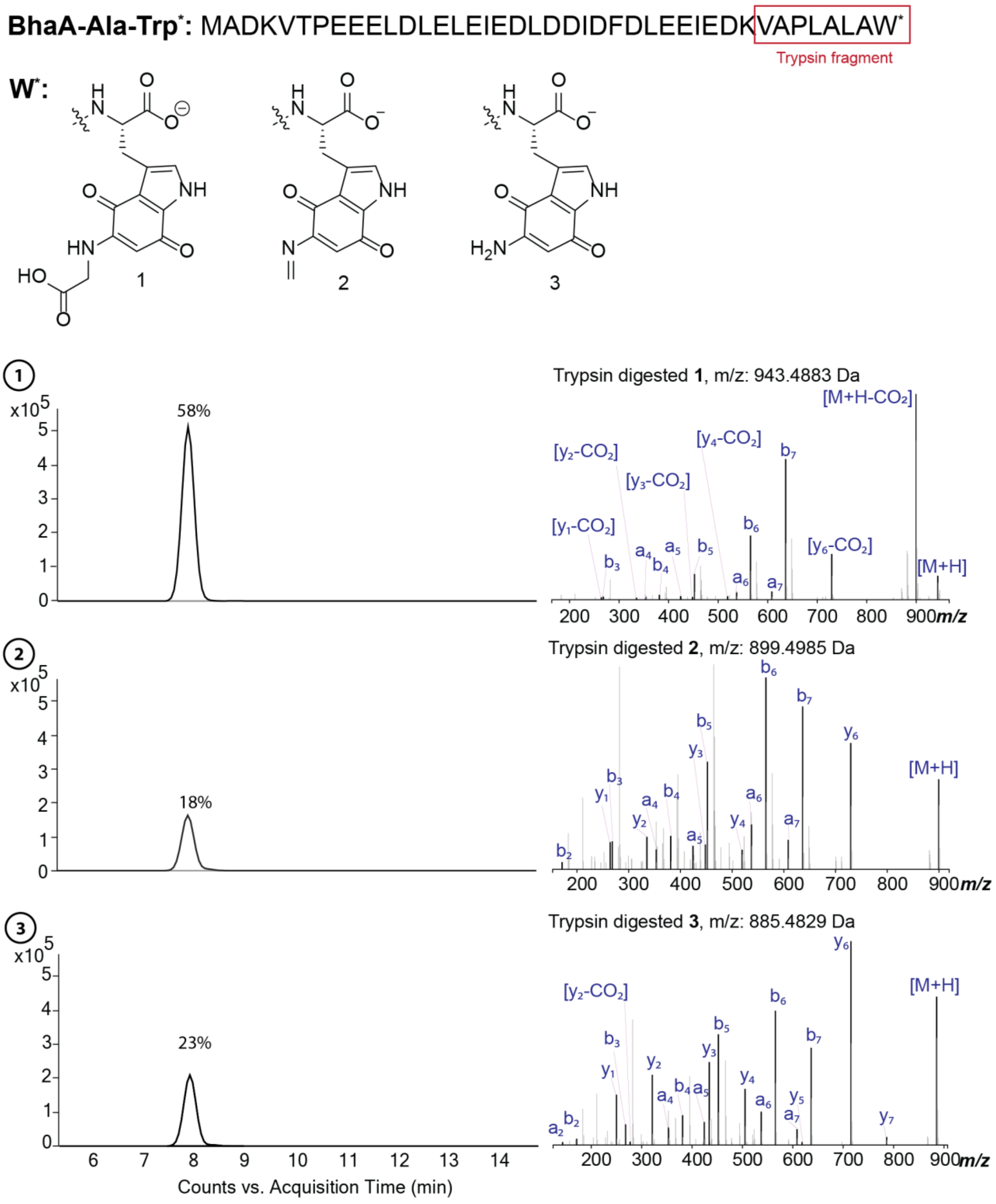


**Figure S2.** ESI-MS/MS analysis of the products of the co-expression of $BhaC_1$ and $BhaB_5$. The mixture of products **1**, **2**, and **3** were digested with trypsin and subjected to LC MS/MS analysis. The major product was the quinone-glycine adduct **1**, followed by compounds **2** and **3** that were produced in similar amounts.

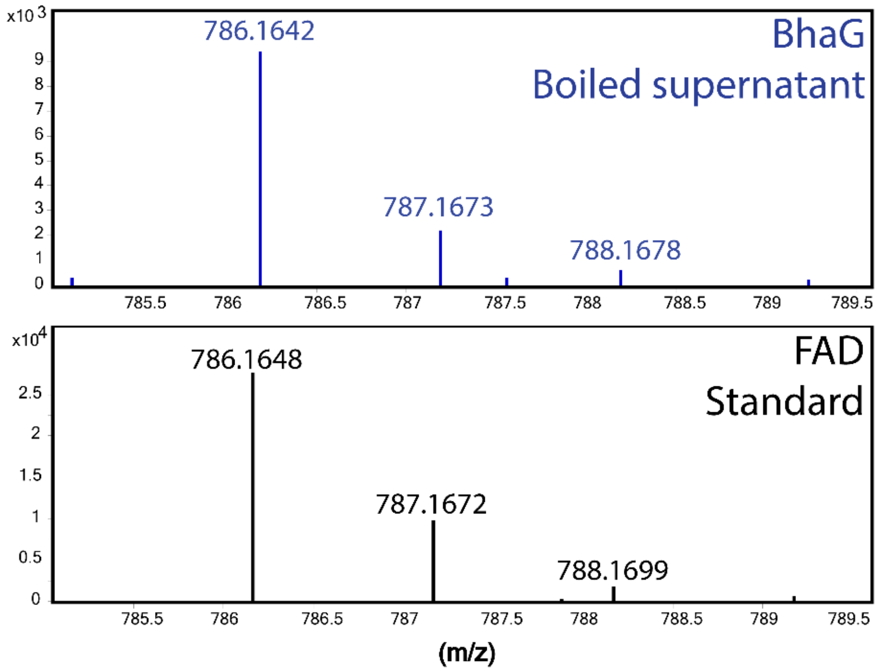


**Figure S3.** BhaG co-purifies with flavin adenine dinucleotide (FAD) as a cofactor. An aliquot of the His-tagged purified BhaG was boiled and the supernatant was submitted for LC-MS analysis and compared with FAD standard.

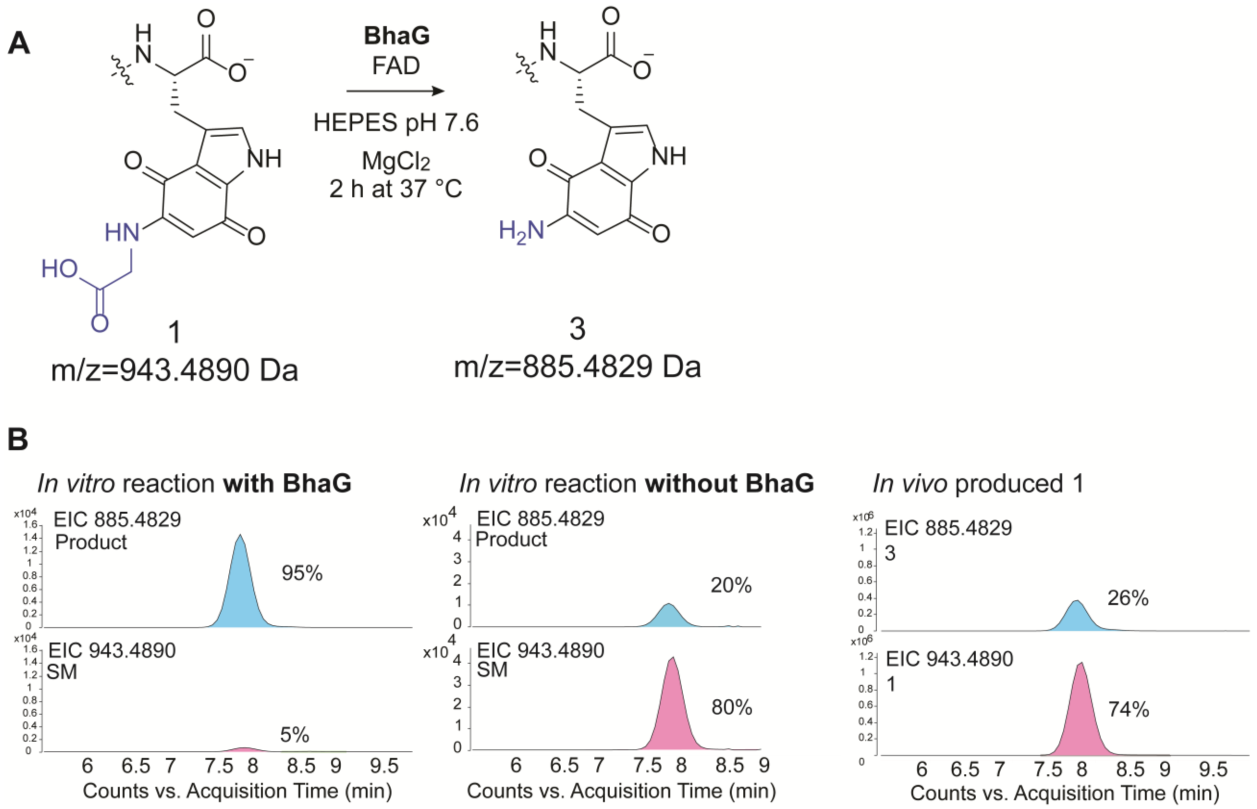


**Figure S4.** In vitro reactions with BhaG. (A) Reaction catalyzed by BhaG. (B) LC-MSMS analysis of a peptide mixture enriched in the glycine-quinone adduct **1** that was reacted with BhaG under the specified reaction conditions. The mixture was then digested with trypsin and analyzed by LC-MS/MS Extracted Ion Chromatography (EIC) showing that incubation with BhaG results in conversion of **1** into aminoquinone **3** (left panel), whereas incubation without BhaG does not result in conversion to aminoquinone **3** (middle panel) beyond background reaction (right panel), which was probably the source of **3** seen in a previous study.[2] SM = starting material. EIC, extracted ion chromatogram.

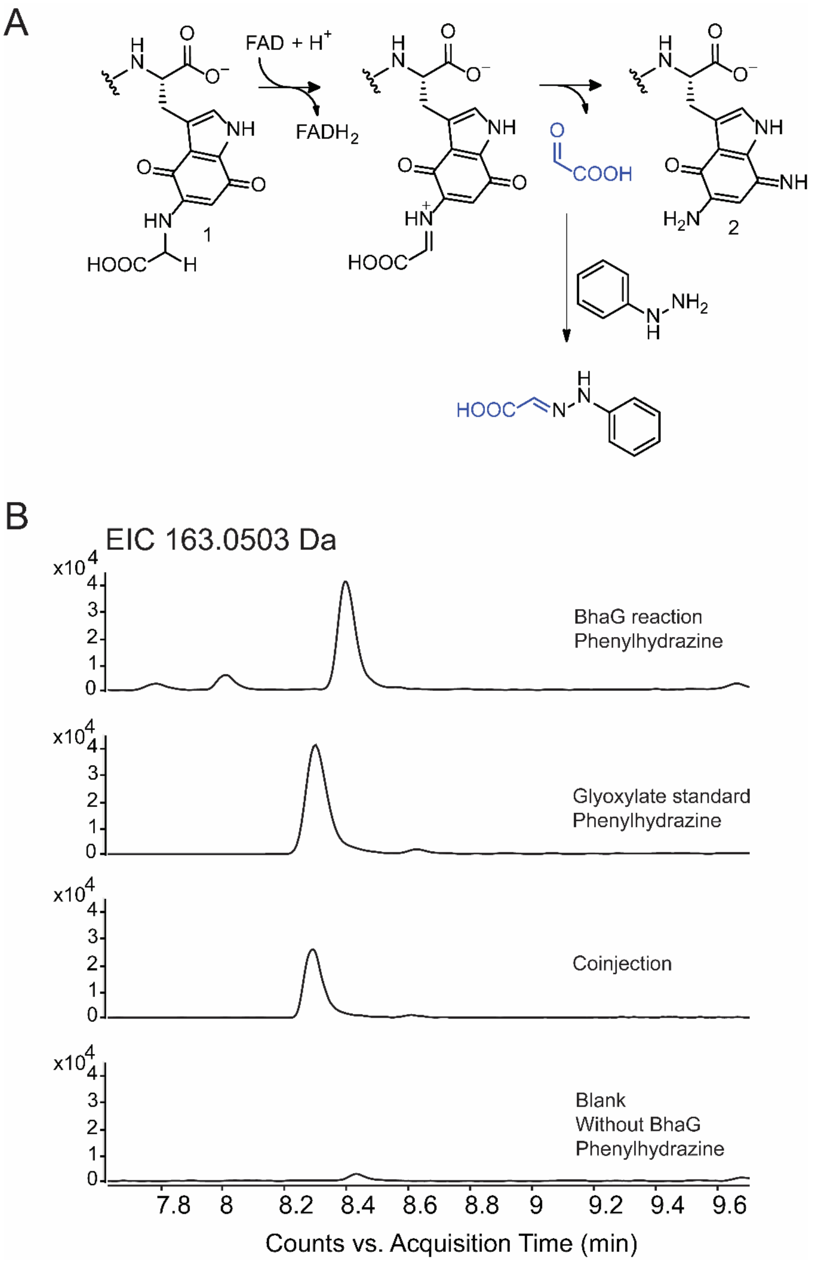


**Figure S5.** Glyoxylate derivatization. (**A**) Reaction mechanism of the BhaG catalyzed reaction. First, hydride transfer from glycine-quinone adduct **1** to FAD generates an imine intermediate. Then, hydration and elimination releases glyoxylic acid and the product aminoquinone **2**. Released glyoxylic acid can be derivatized with phenylhydrazine forming the corresponding phenylhydrazone derivative. Molecular oxygen presumably regenerates the oxidized FAD cofactor. (**B**) LC-MS analysis of the BhaG reaction products derivatized with phenylhydrazine. EICs monitoring the hydrazone adduct. Top trace: product of the in vitro BhaG reaction treated with phenylhydrazine. 2nd trace: glyoxylate authentic standard treated with phenylhydrazine. 3rd trace: coinjection of the glyoxylate authentic standard treated with phenylhydrazine and the BhaG product trated with phenylhydrazine. Bottom trace: Reaction mixture without enzyme treated with phenylhydrazine.

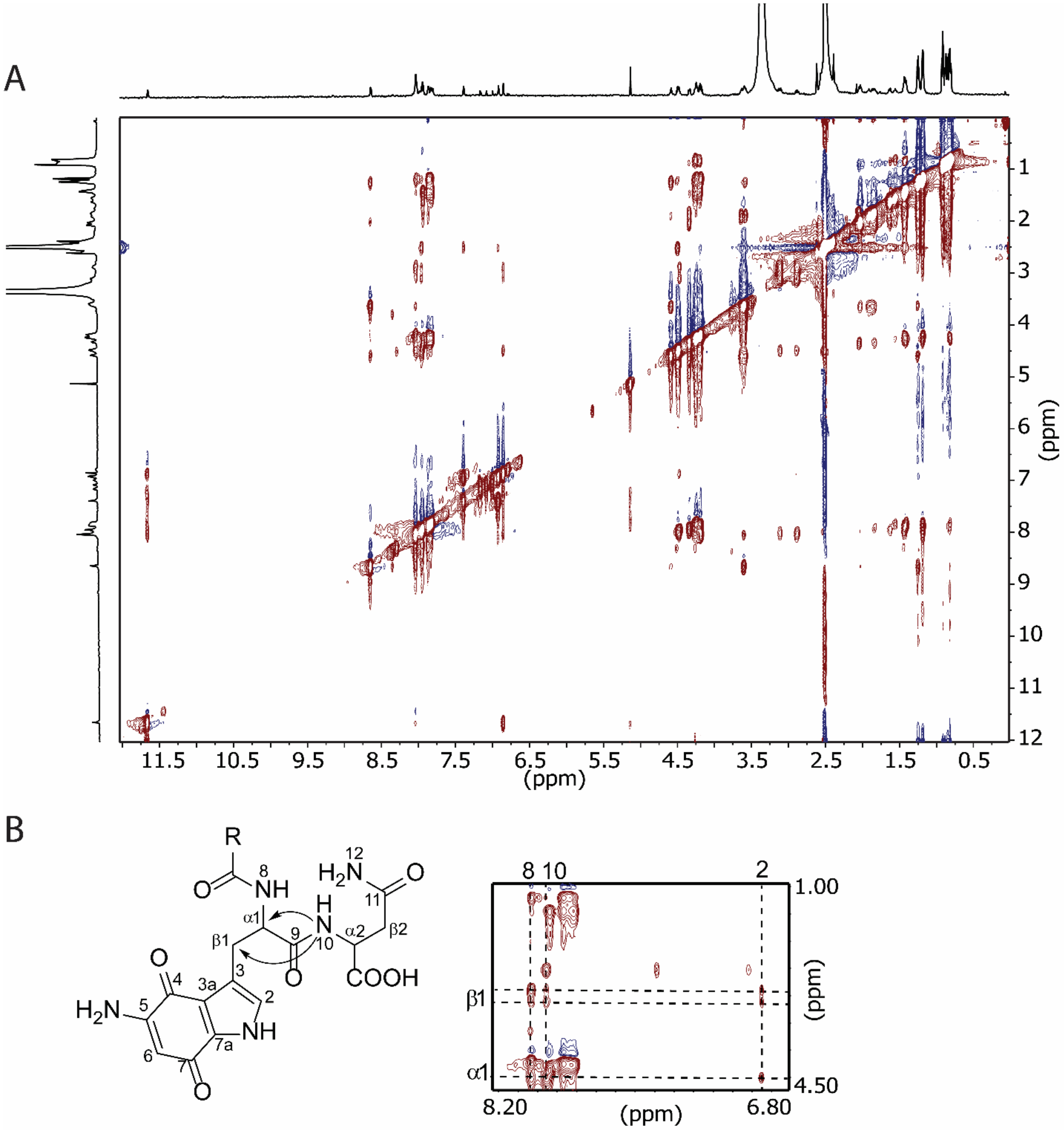
A
11.5
10.5
9.5
8.5
7.5
6.5
5.5
4.5
3.5
2.5
1.5
0.5
(ppm)
B
R
O
8
NH
$H_2N$
12
11
O
H
N
10
α1
β1
9
α2
β2
O
COOH
3
3a
4
5
$H_2N$
6
7
7a
2
NH
8 10
2
1.00
β1
α1
4.50
8.20
(ppm)
6.80

C

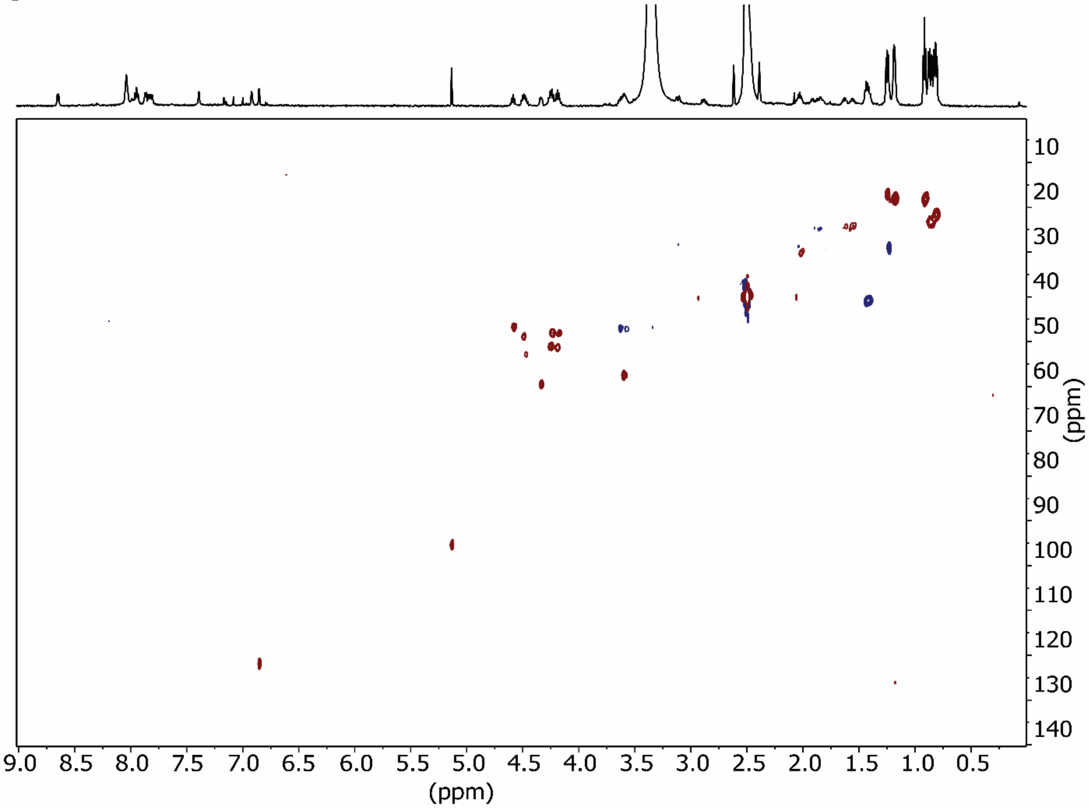

D

| C/H | δH | δC |
|---|---|---|
| α1 | 4.45 | |
| β1 | 3.10, 2.87 | 28.34 |
| 1 | 11.65 | |
| 2 | 6.85 | 121.86 |
| 3 | | n.o. |
| 3a | | 119.98 |
| 4 | | 178.85 |
| 5 | | n.o. |
| 6 | 5.13 | 95.57 |
| 7 | | n.o. |
| 7a | | 131.15 |
| 8, Trp NH | 8.06 | |
| 9, Trp C=O | | 172.46 |
| α2 | 4.48 | |
| β2 | 2.49 | |
| 11 | | 172.14 |
| 12 | 7.38, 6.90 | |
| 10, Asn NH | 7.95 | |

**Figure S6.** Nuclear Magnetic Resonance (NMR) spectra of BhaB$_4$ product **4**. (**A**) NOESY spectrum of trypsin-digested **4**. (**B**) Diagnostic NOESY correlations are indicated with black arrows. (**C**) HSQC spectrum of trypsin-digested **4**. (**D**) $^{1}$H and $^{13}$C NMR assignments for trypsin digested **4**, n.o. indicates not observed.

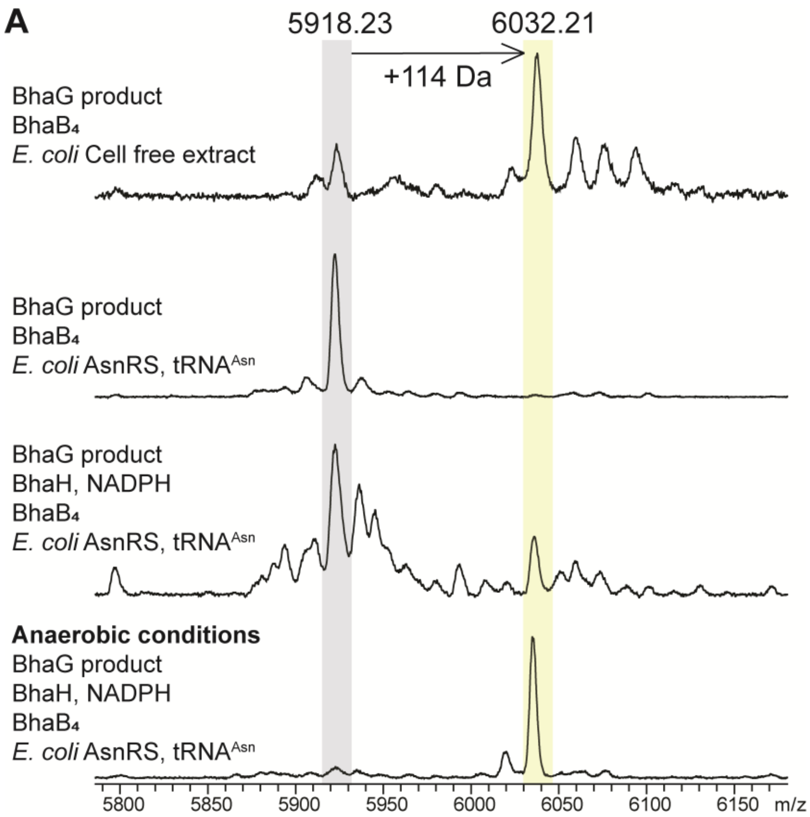
A
5918.23
6032.21
+114 Da
BhaG product
BhaB₄
E. coli Cell free extract
BhaG product
BhaB₄
E. coli AsnRS, tRNAAsn
BhaG product
BhaH, NADPH
BhaB₄
E. coli AsnRS, tRNAAsn
Anaerobic conditions
BhaG product
BhaH, NADPH
BhaB₄
E. coli AsnRS, tRNAAsn
5800
5850
5900
5950
6000
6050
6100
6150
m/z

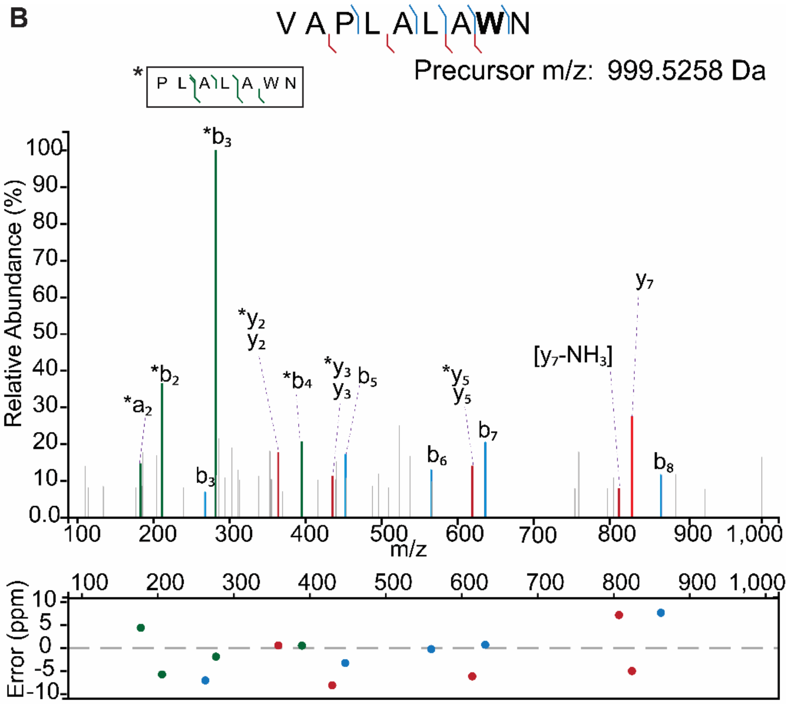


**VAPLALAWN**

| Fragment Type | Experimental Mz | Theoretical Mz | Mass Error (ppm) |
|---|---|---|---|
| $b_3$ | 268.1637 | 268.1656 | -6.9676 |
| $y_2$ | 364.1254 | 364.1252 | 0.601 |
| $y_3$ | 435.1588 | 435.1623 | -8.0318 |
| $b_5$ | 452.2853 | 452.2867 | -3.1893 |
| $b_6$ | 565.3707 | 565.3708 | -0.1812 |
| $y_5$ | 619.2797 | 619.2835 | -6.0863 |
| $b_7$ | 636.4084 | 636.4079 | 0.7598 |
| $y_7$ (-$NH_3$) | 812.3996 | 812.3937 | 7.2083 |
| $y_7$ | 829.4162 | 829.4203 | -4.935 |
| $b_8$ | 867.479 | 867.4723 | 7.6931 |

***PLALAWN**

| Fragment Type | Experimental Mz | Theoretical Mz | Mass Error (ppm) |
|---|---|---|---|
| *$a_2$ | 183.15 | 183.1492 | 4.4453 |
| *$b_2$ | 211.1429 | 211.1441 | -5.6855 |
| *$b_3$ | 282.1807 | 282.1812 | -1.8232 |
| *$y_2$ | 364.1254 | 364.1252 | 0.601 |
| *$b_4$ | 395.2655 | 395.2653 | 0.5706 |
| *$y_3$ | 435.1588 | 435.1623 | -8.0318 |
| *$y_5$ | 619.2797 | 619.2835 | -6.0863 |

**Figure S7**. (**A**) MALDI-TOF mass spectra of in vitro reactions containing BhaG product (aminoquinone intermediate **3**), and $BhaB_4$ using *E. coli* cell-free extract (top panel), using *E. coli* AsnRS and $tRNA^{Asn}$ (second panel), including BhaH in addition to *E. coli* AsnRS and $tRNA^{Asn}$ (third panel), and using anaerobic conditions with BhaH in addition to *E. coli* AsnRS and $tRNA^{Asn}$ (bottom panel). (**B**) ESI-MSMS analysis of in vitro produced unlabeled $BhaB_4$ product **4**. Cleavage of the amide bond at AP during LC-MS/MS analysis produces a shorter fragment PLALAWN with the corresponding b and y ions. This fragmentation is likely due to the Pro effect.[3-6] A graph of the ppm errors for each identified ion is shown.[7]

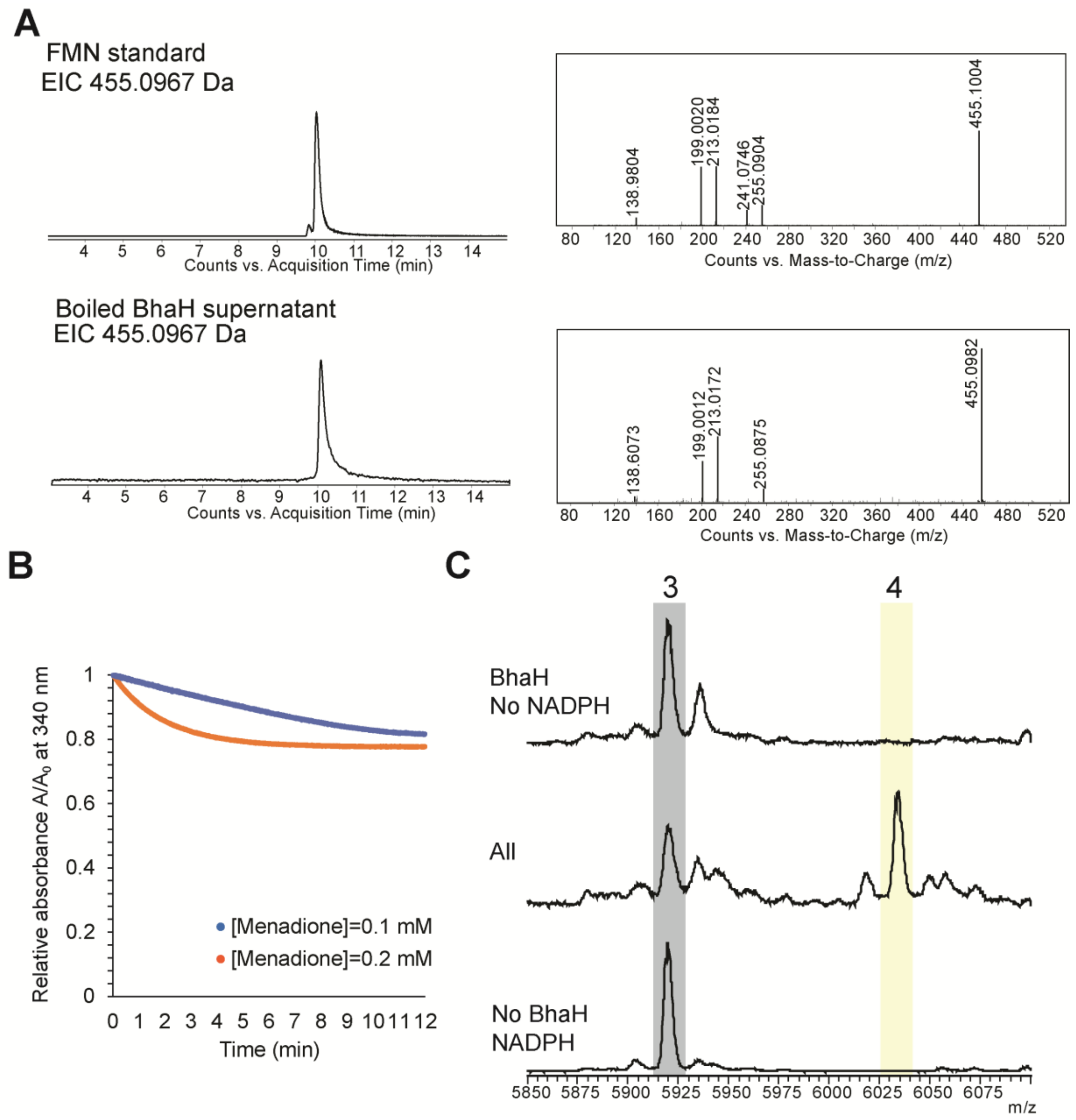


**Figure S8**. (**A**) BhaH co-purifies with flavin mononucleotide (FMN) as a cofactor. (**B**) BhaH activity assay using menadione as a substrate under anaerobic conditions. During the catalytic cycle, the bound cofactor, FMN, is reduced by NADPH. Then, the resulting $FMNH_2$ reduces menadione regenerating FMN. (**C**) $BhaB_4$ activity was tested in the absence of BhaH or NADPH, no conversion to product **4** was observed. However, when both, BhaH and NADPH were added to the mixture, conversion to product **4** is observed.

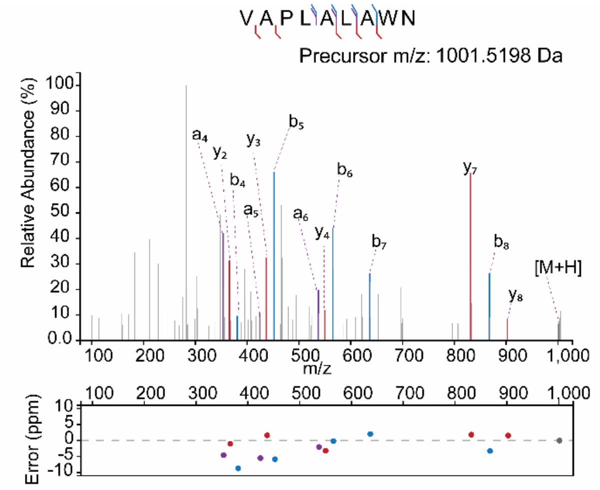


| Fragment Type | Experimental Mz | Theoretical Mz | Mass Error (ppm) |
|---|---|---|---|
| a4 | 353.2531 | 353.2547 | -4.5685 |
| y2 | 366.1188 | 366.1192 | -1.0411 |
| b4 | 381.2463 | 381.2496 | -8.7304 |
| a5 | 424.2895 | 424.2918 | -5.4864 |
| y3 | 437.157 | 437.1563 | 1.6124 |
| b5 | 452.2841 | 452.2867 | -5.8425 |
| a6 | 537.3748 | 537.3759 | -2.0244 |
| y4 | 550.2386 | 550.2404 | -3.1898 |
| b6 | 565.3707 | 565.3708 | -0.1812 |
| b7 | 636.4092 | 636.4079 | 2.0168 |
| y7 | 831.4158 | 831.4143 | 1.8124 |
| b8 | 867.4695 | 867.4723 | -3.2583 |
| y8 | 902.4528 | 902.4514 | 1.5434 |
| M+H | 1001.5198 | 1001.5198 | -0.0211 |

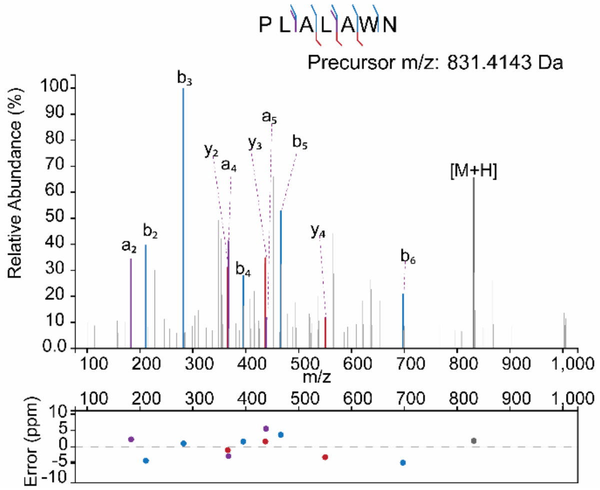


| Fragment Type | Experimental Mz | Theoretical Mz | Mass Error (ppm) |
|---|---|---|---|
| $a_2$ | 183.1496 | 183.1492 | 2.2613 |
| $b_2$ | 211.1432 | 211.1441 | -4.2647 |
| $b_3$ | 282.1815 | 282.1812 | 1.0119 |
| $y_2$ | 366.1188 | 366.1192 | -1.0411 |
| $a_4$ | 367.2693 | 367.2704 | -2.8857 |
| $b_4$ | 395.2659 | 395.2653 | 1.5826 |
| $y_3$ | 437.157 | 437.1563 | 1.6124 |
| $a_5$ | 438.3099 | 438.3075 | 5.5353 |
| $b_5$ | 466.3041 | 466.3024 | 3.6704 |
| $y_4$ | 550.2386 | 550.2404 | -3.1898 |
| $b_6$ | 697.3634 | 697.3668 | -4.8733 |
| M+H | 831.4158 | 831.4143 | 1.8124 |

**Figure S9**. LC-MS/MS analysis of $BhaB_4$ product obtained from *in vitro* reaction using labeled L-$^{15}N_2$-Asn, $tRNA^{Asn}$, AsnRS, and BhaH . Cleavage N-terminal to the Pro produces a labeled PLALA**W**N with the corresponding b and y ions, likely due to the Pro effect.[3-6] A graph of the ppm errors for each identified ion is shown.[7] **W** = modified Trp in **4** (see text).

**A**

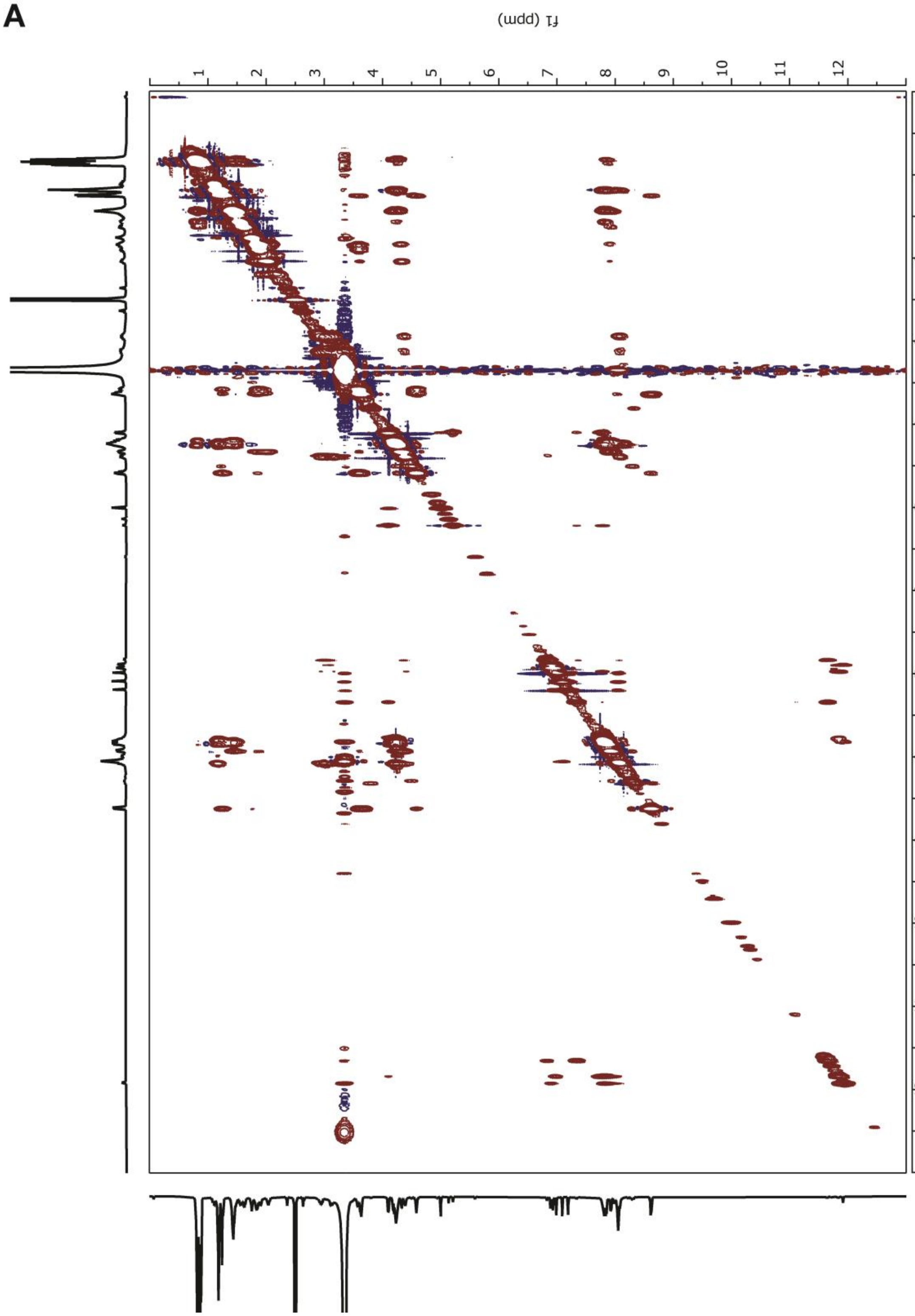

**B**

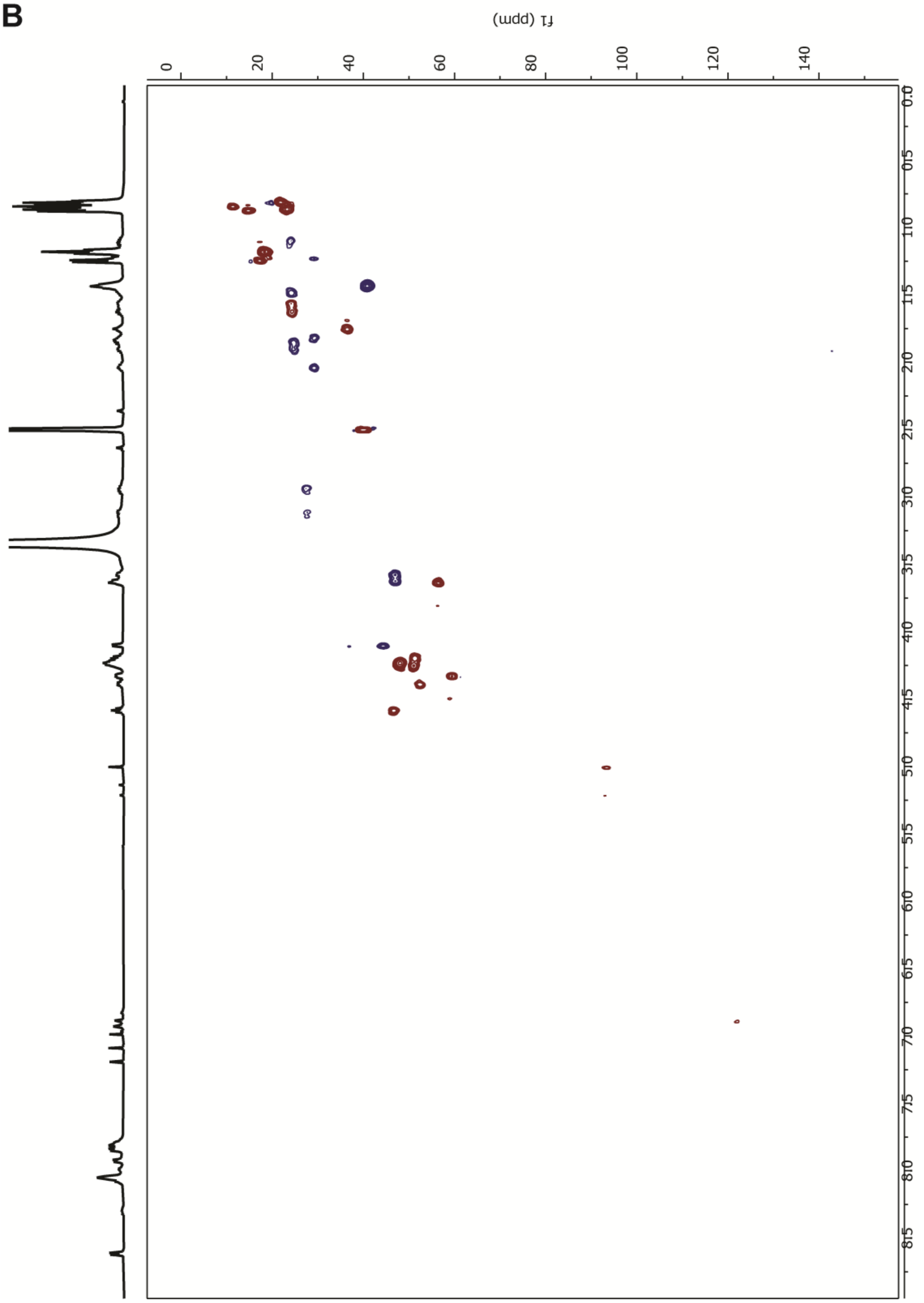

C

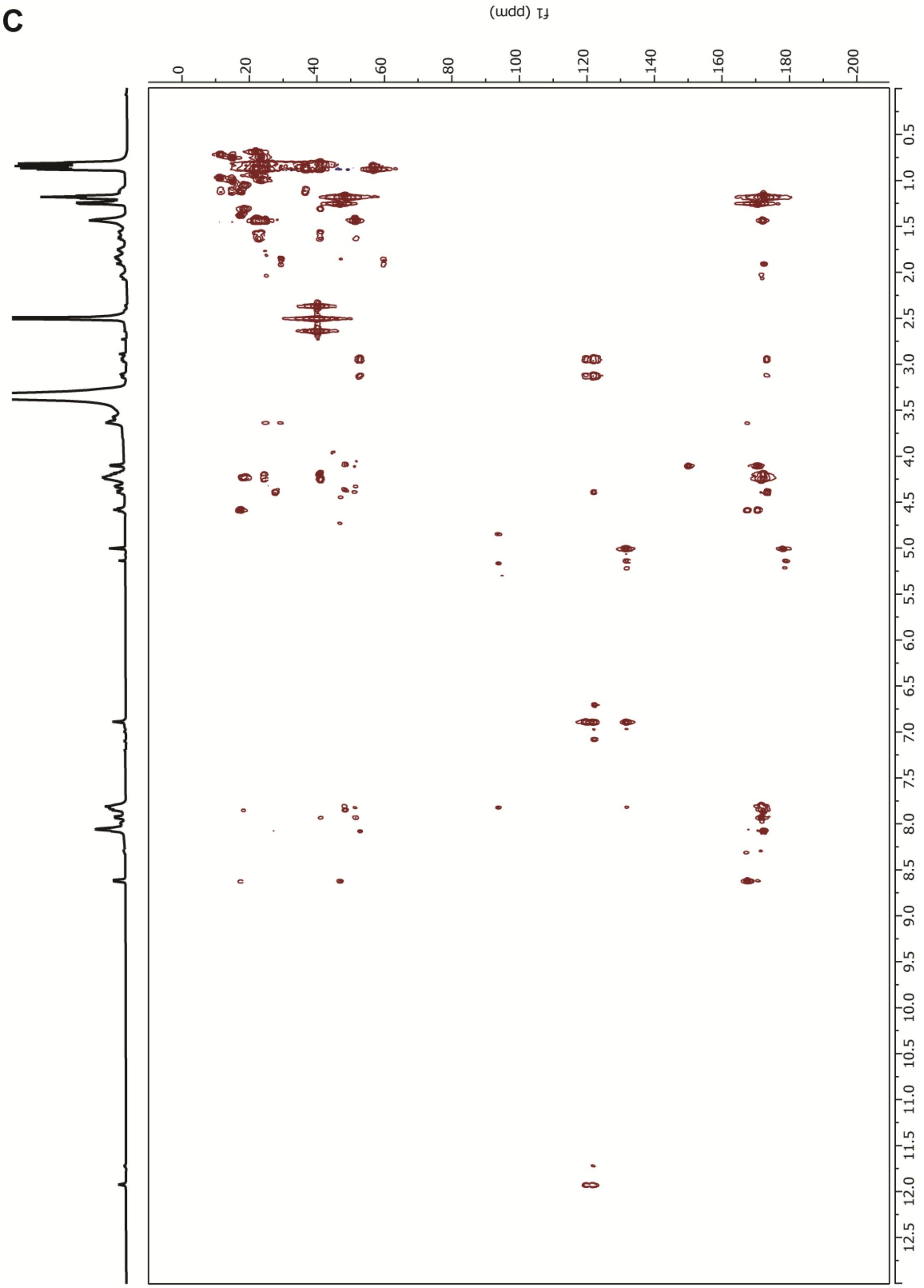

**D**

R=IAPLALA

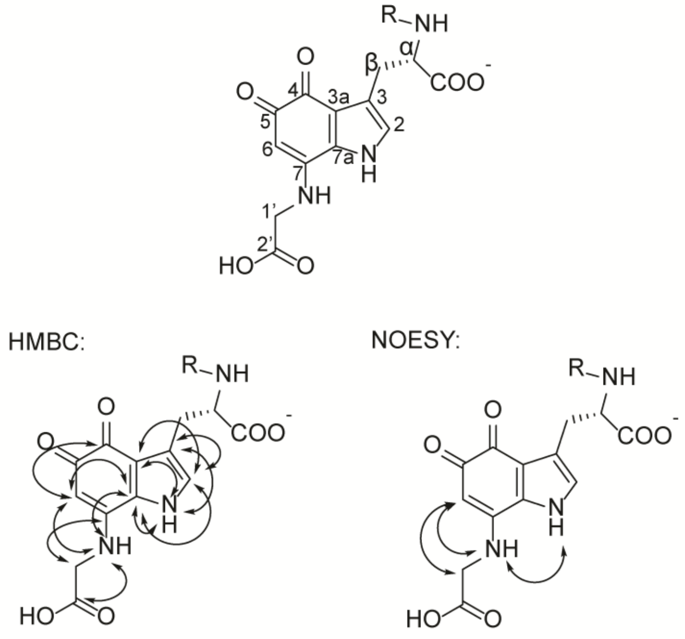


| Position | δ $^{13}$C | δ $^{1}$H |
|---|---|---|
| 2 | 122.4 | 6.89 |
| 3 | 119.7 | - |
| 3a | 122.1 | - |
| NH (Im ring) | n.a. | 11.92 |
| 4 | 178.2 | - |
| 5 | Not observed | - |
| 6 | 93.9 | 5.00 |
| 7 | 150.2 | - |
| 7a | 131.9 | - |
| 1' (Gly-$CH_2$) | 44.7 | 4.10 |
| 2' (Gly-COOH) | 170.6 | - |
| NH (Gly) | n.a. | 7.82 |
| α | 52.1 | 4.39 |
| β | 27.1 | 3.14, 2.94 |

**E**

R=AmmA*

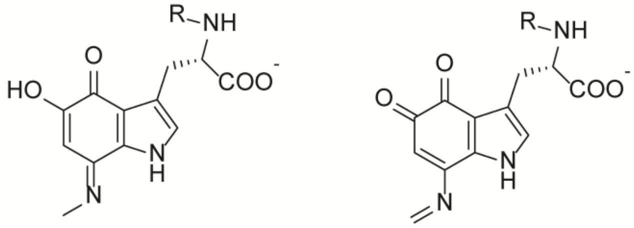

**2'** **2''**

**Figure S10**. NMR spectra of $AmmB_3$ product, **5**. (**A**) NOESY spectrum of trypsin-digested **5**. (**B**) HSQC spectrum of trypsin-digested **5**. (**C**) HMBC spectrum of trypsin-digested **5**. (**D**) $^{1}$H and $^{13}$C NMR assignments for trypsin digested **5**. Diagnostic HMBC and NOESY correlations are indicated with black arrows. (**E**) Imine product **2''** or its possible tautomer **2'** formed by $AmmB_3$ in *E. coli* based on MS data reported previously.[2]

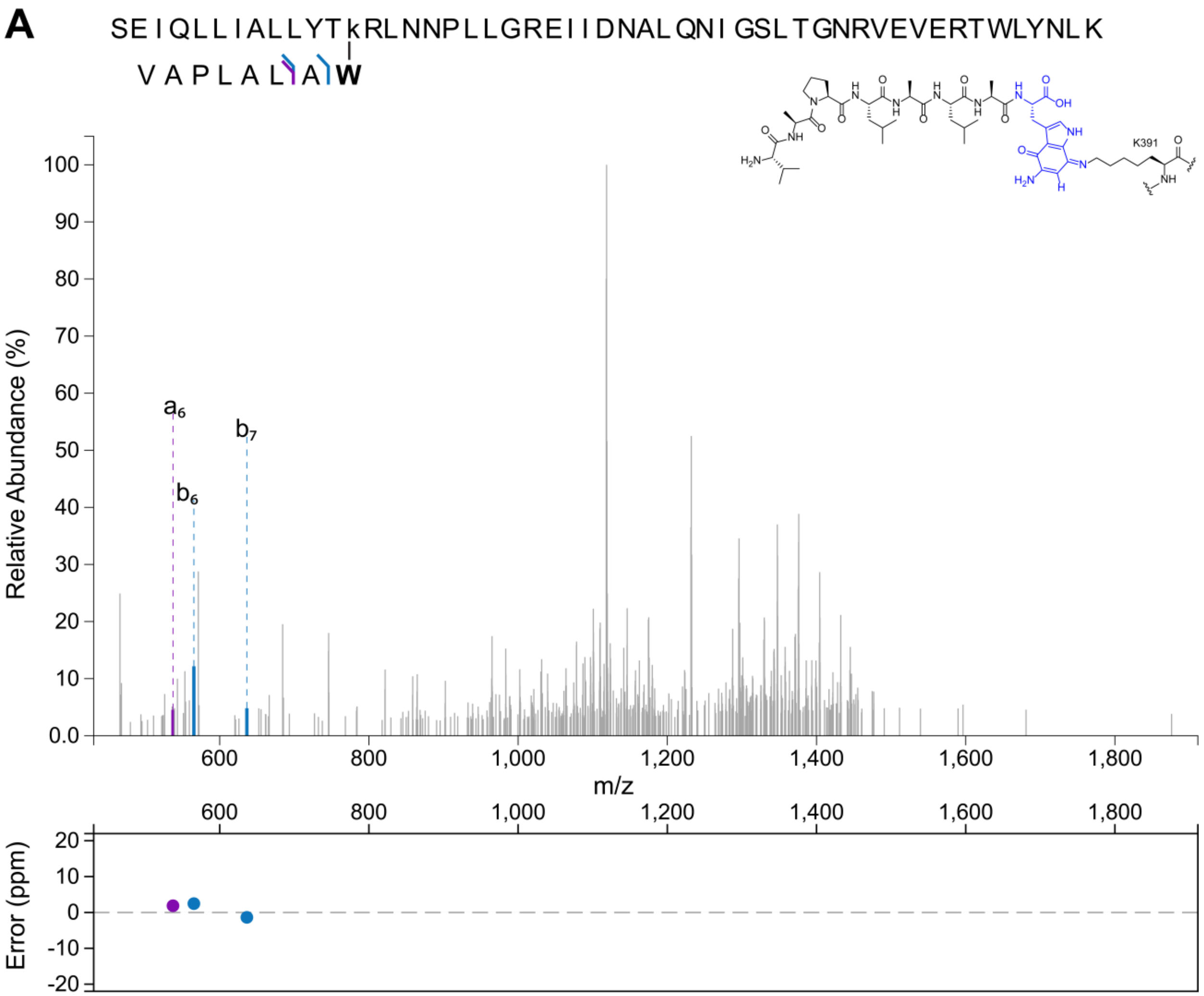
A
SEIQLLIALLYTkRLNNPLLGREIIDNALQNIGSLTGNRVEVERTWLYNLK
VAPLALAW
K391
Relative Abundance (%)
a6
b6
b7
m/z
Error (ppm)

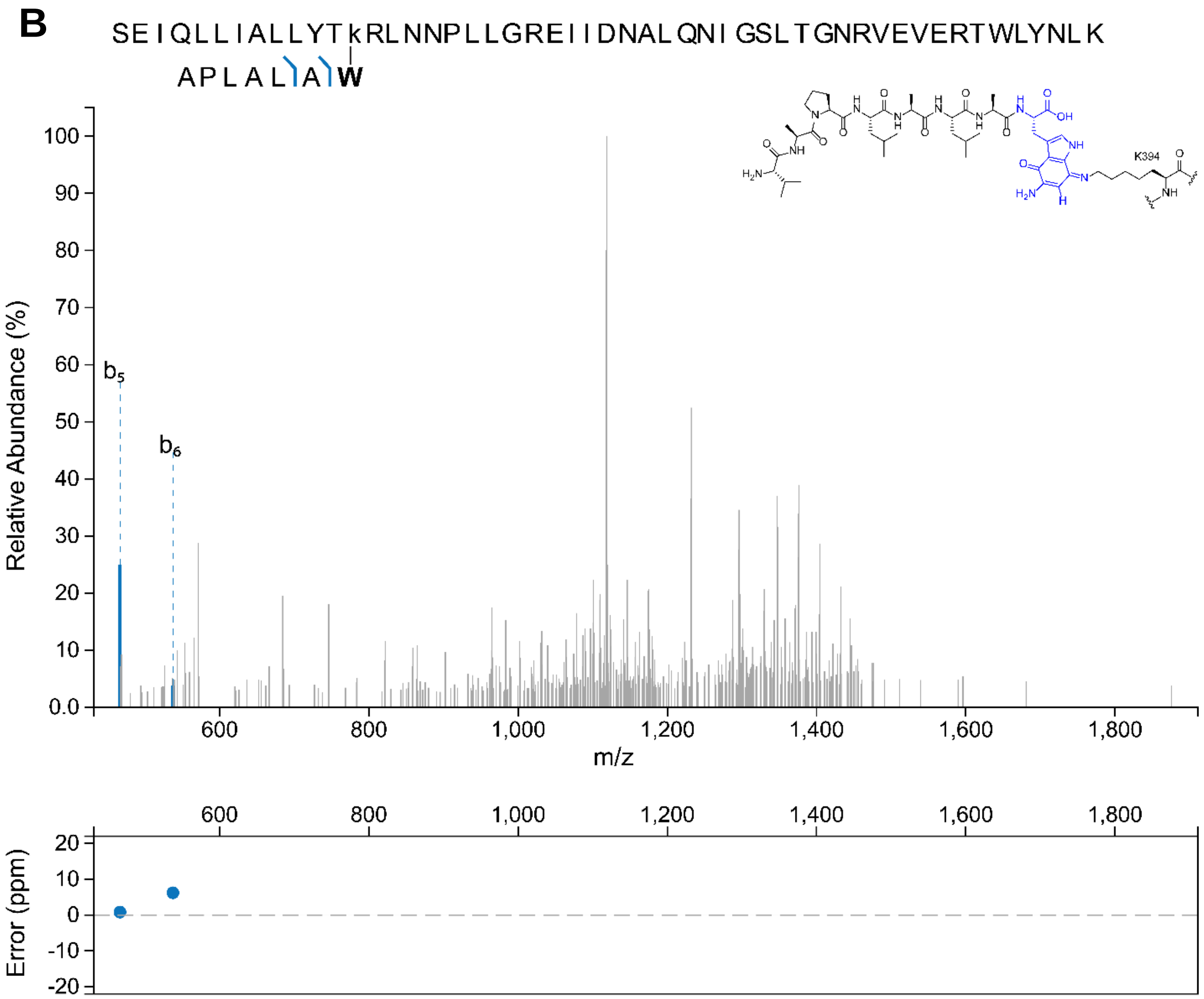
B
SEIQLLIALLYTkRLNNPLLGREIIDNALQNIGSLTGNRVEVERTWLYNLK
APLALAW
b5
b6
Relative Abundance (%)
m/z
Error (ppm)
K394

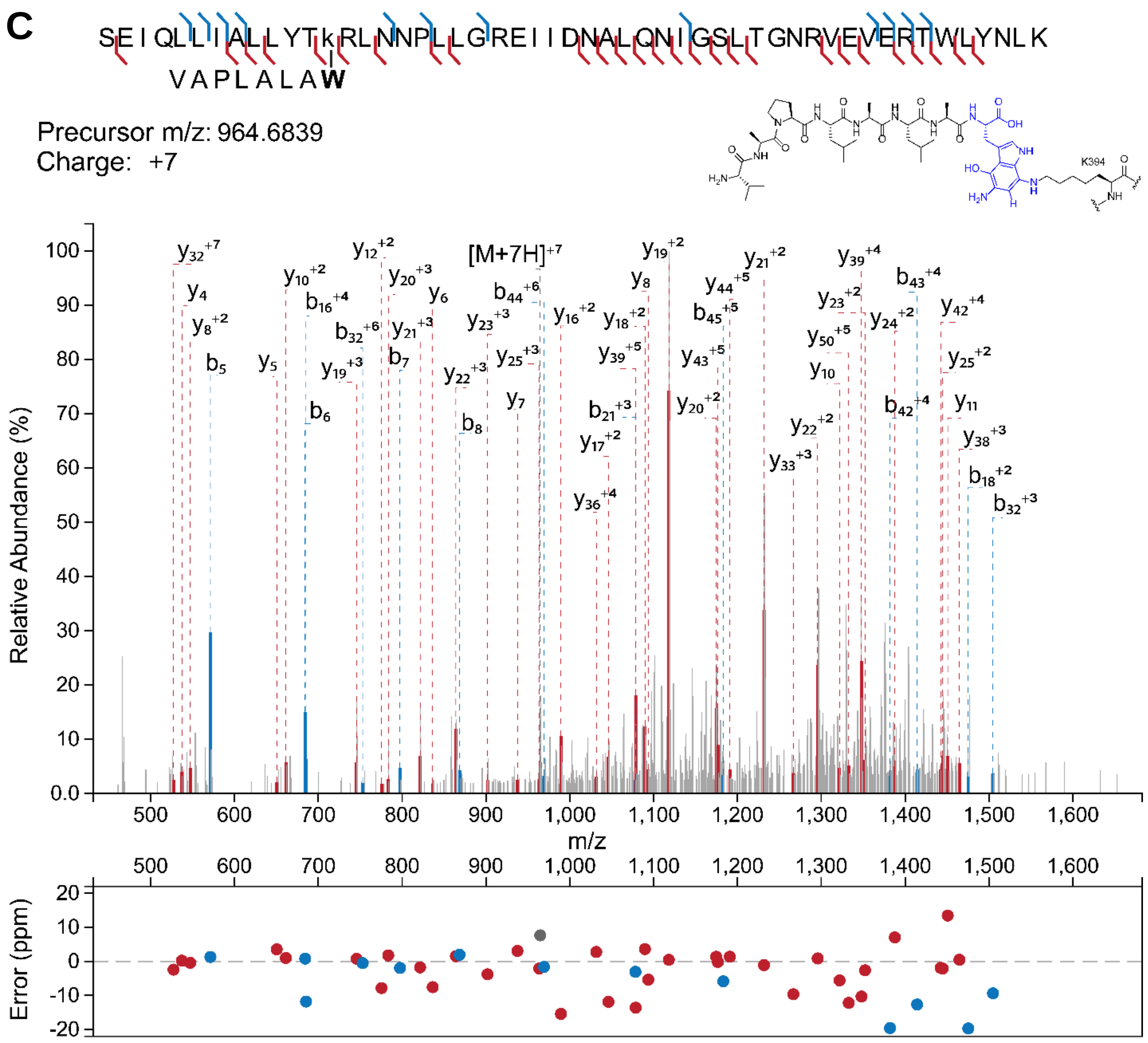
C
SEIQLLIALLYTkRLNNPLLGREIIDNALQNIGSLTGNRVEVERTWLYNLK
VAPLALAW
Precursor m/z: 964.6839
Charge: +7
K394
[M+7H]+7
Relative Abundance (%)
m/z
Error (ppm)

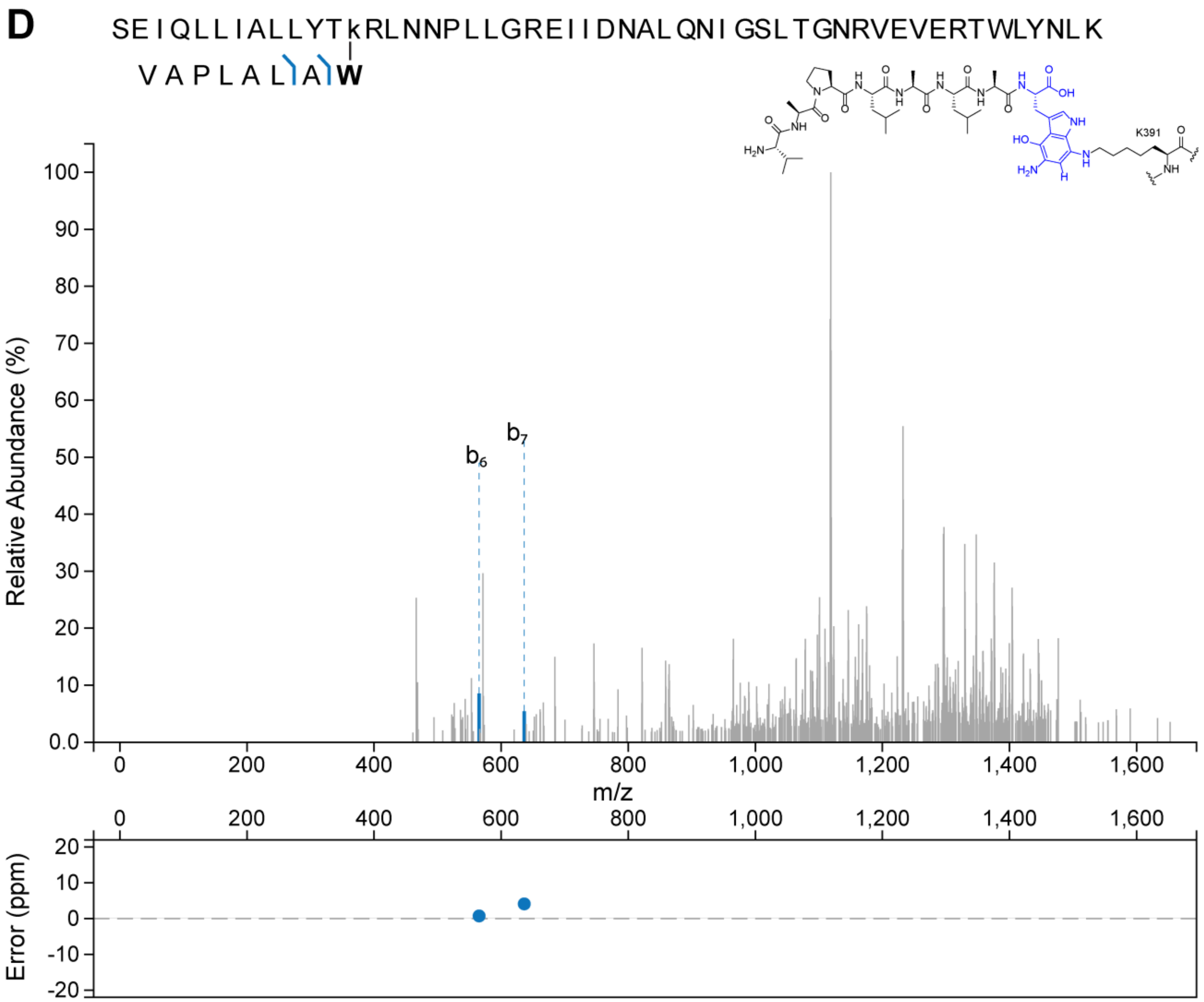
D
SEIQLLIALLYTkRLNNPLLGREIIDNALQNIGSLTGNRVEVERTWLYNLK
VAPLALIAW
K391
Relative Abundance (%)
b6
b7
m/z
Error (ppm)

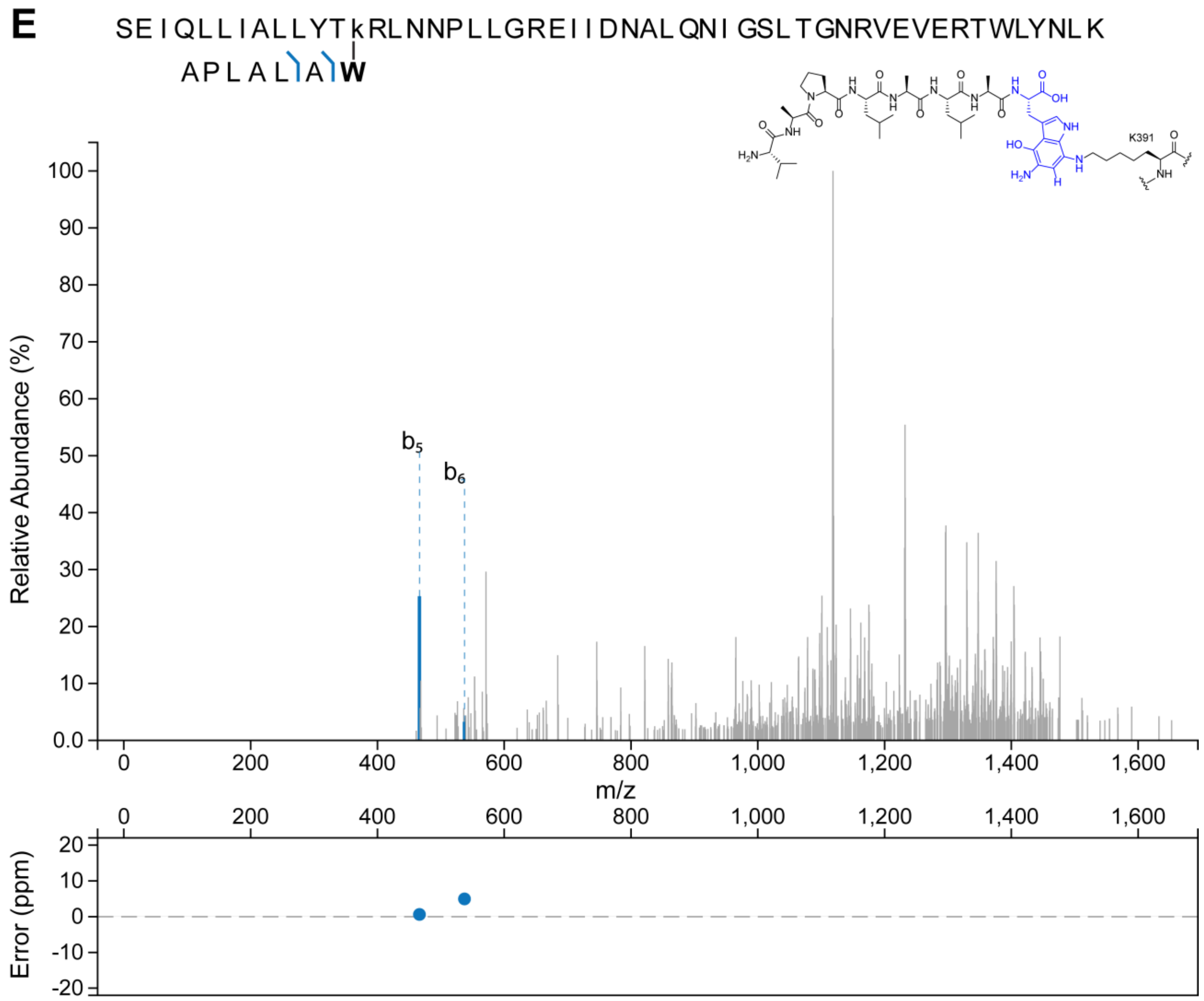


**Figure S11.** $BhaC_1$ crosslink analysis with intermediate **3**. The crosslink is drawn at C7 of the indole, but it could also be at C4. The tandem MS spectrum in panels A and B is the same as Figure 4B, main text, but additional peaks not annotated in Figure 4B are annotated here. (A) Assignment of additional fragments to the $a_6$, $b_6$ and $b_7$ ions of the linked modified BhaA-AW peptide after LysC digestion. (B) Cleavage of the linked peptide at Val-Ala produces fragments corresponding to $b_5$ and $b_6$ ions of the crosslinked BhaA peptide. (C) ESI-MS/MS data on a $BhaC_1$ fragment that is crosslinked with intermediate **3** in the reduced hydroquinone form. (D) $b_6$ and $b_7$ ions of the BhaA peptide in the reduced hydroquinone form linked to the $BhaC_1$ fragment peptide produced by LysC digestion. (E) Cleavage of the linked BhaA peptide in the reduced hydroquinone form at Val-Ala produces fragments corresponding to $b_5$ and $b_6$ ions of the crosslinked BhaA.

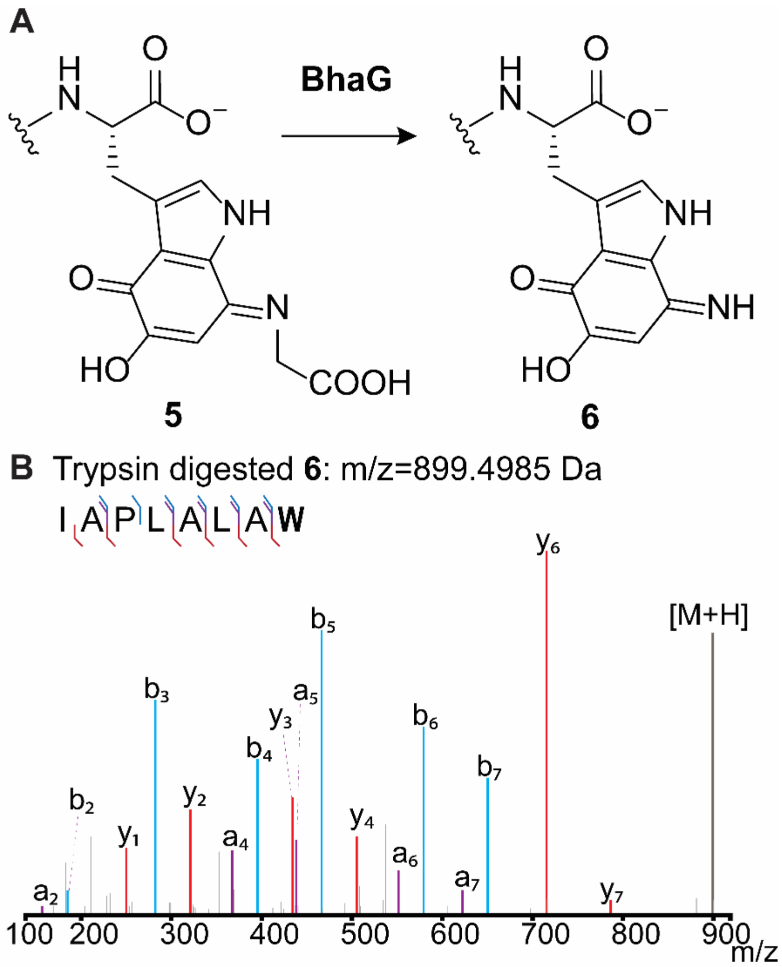


C

| Fragment Type | Experimental Mz | Theoretical Mz | Mass Error (ppm) |
|---|---|---|---|
| $a_1$ | 157.1342 | 157.1335 | 4.2267 |
| $b_2$ | 185.1292 | 185.1285 | 4.0487 |
| $y_1$ | 250.0832 | 250.0823 | 3.7941 |
| $b_3$ | 282.1832 | 282.1812 | 7.0364 |
| $y_2$ | 321.1209 | 321.1194 | 4.7797 |
| $a_4$ | 367.2723 | 367.2704 | 5.2826 |
| $b_4$ | 395.2674 | 395.2653 | 5.3775 |
| $y_3$ | 434.2057 | 434.2034 | 5.2391 |
| $a_5$ | 438.3099 | 438.3075 | 5.5353 |
| $b_5$ | 466.3057 | 466.3024 | 7.1017 |
| $y_4$ | 505.2430 | 505.2405 | 4.8706 |
| $a_6$ | 551.3943 | 551.3915 | 5.0167 |
| $b_6$ | 579.3901 | 579.3864 | 6.3024 |
| $a_7$ | 622.4318 | 622.4286 | 5.0643 |
| $b_7$ | 650.4278 | 650.4236 | 6.5150 |
| $y_6$ | 715.3826 | 715.3774 | 7.3204 |
| $y_7$ | 786.4181 | 786.4145 | 4.6068 |
| M+H | 899.5056 | 899.4985 | 7.8520 |

**Figure S12**. (**A**) Aminoquinone, **6**, is formed upon addition of BhaG to the co-expression of AmmA*-Trp with $BhaC_1$ ($AmmC_1$ ortholog) and $AmmB_3$. (**B**) LC-MS/MS analysis of the trypsin digested fragment **6**. **W** = modified Trp.

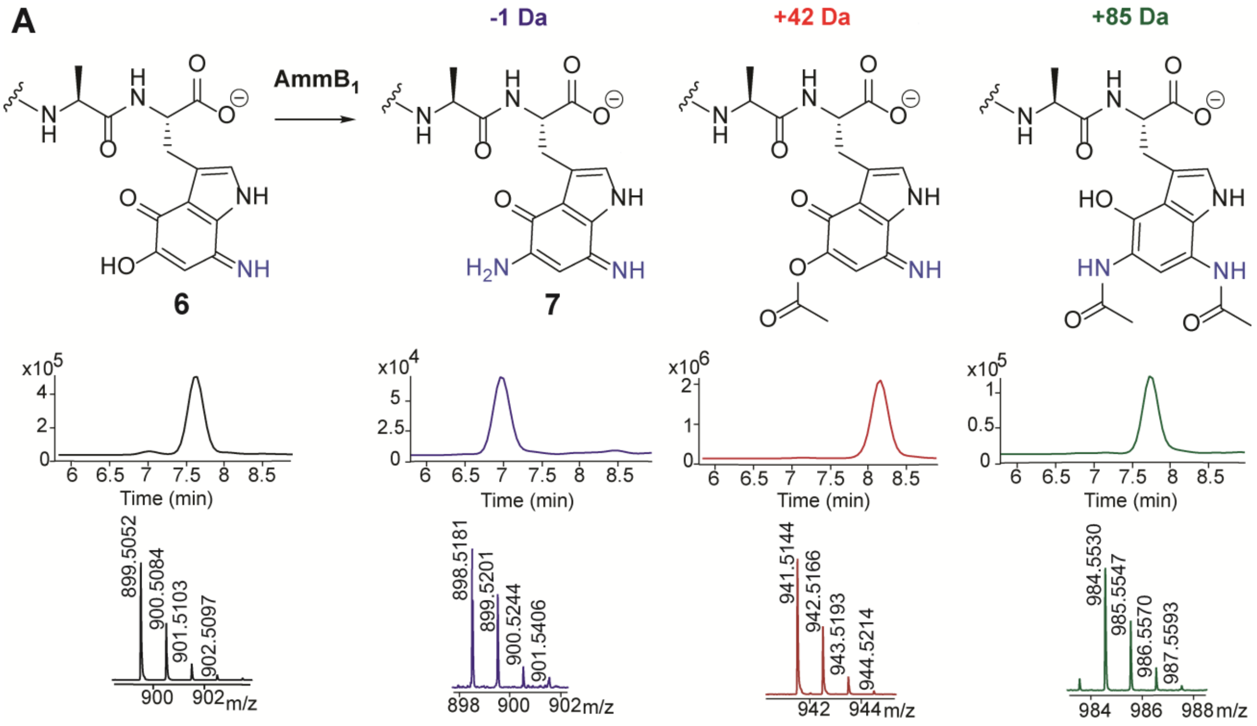


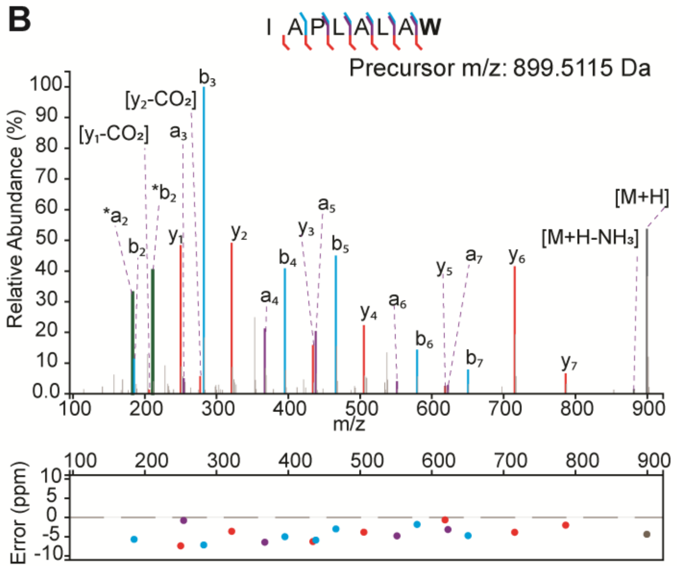


| Fragment Type | Experimental Mz | Theoretical Mz | Mass Error (ppm) |
|---|---|---|---|
| *$a_2$ | 183.1474 | 183.1492 | -9.7508 |
| *$b_2$ | 211.1425 | 211.1441 | -7.58 |
| $a_2$ | 157.1319 | 157.1335 | -10.4106 |
| $b_2$ | 185.1274 | 185.1285 | -5.6743 |
| $y_1$ ($-CO_2$) | 206.1042 | 206.1054 | -5.9286 |
| $y_1$ | 250.0934 | 250.0953 | -7.4018 |
| $a_3$ | 254.1861 | 254.1863 | -0.7862 |
| $y_2$ ($-CO_2$) | 277.1403 | 277.1425 | -8.0677 |
| $b_3$ | 282.1792 | 282.1812 | -7.1389 |
| $y_2$ | 321.1312 | 321.1324 | -3.6283 |
| $a_4$ | 367.268 | 367.2704 | -6.4254 |
| $b_4$ | 395.2633 | 395.2653 | -4.9953 |
| $y_3$ | 434.2137 | 434.2164 | -6.276 |
| $a_5$ | 438.3049 | 438.3075 | -5.8722 |
| $b_5$ | 466.301 | 466.3024 | -2.9776 |
| $y_4$ | 505.2516 | 505.2535 | -3.838 |
| $a_6$ | 551.3889 | 551.3915 | -4.7767 |
| $b_6$ | 579.3854 | 579.3864 | -1.8096 |
| $y_5$ | 618.3372 | 618.3376 | -0.6455 |
| $a_7$ | 622.4267 | 622.4286 | -3.1294 |
| $b_7$ | 650.4205 | 650.4236 | -4.7084 |
| $y_6$ | 715.3876 | 715.3904 | -3.8624 |
| $y_7$ | 786.4259 | 786.4275 | -2.0055 |
| M+H ($-NH_3$) | 882.4937 | 882.485 | 9.8721 |
| M+H | 899.5076 | 899.5115 | -4.377 |

**Figure S13**. (**A**) Products of co-expression of AmmB$_1$, AmmA*-Trp, BhaC$_1$ (AmmC$_1$ ortholog), AmmB$_3$, and BhaG. Trypsin digested fragments were generated and analyzed with LC-MS/MS. These fragments contained IAPLALAW with mass shifts of -1 (experimental m/z=898.5181, theoretical m/z=898.5145, error=4.0066 ppm), +42 (experimental m/z=941.5144, theoretical m/z=941.5091, error=5.6292 ppm), and +85 Da (experimental m/z=984.5530, theoretical m/z=984.5513, error=1.7267 ppm), compared to peptide **6**. The monoacetylated product is drawn as residing on O5 but it could also be on N7. (**B**) LC-MS/MS analysis of product **7** obtained from *in vitro* reaction with AmmB$_1$ using labeled L-$^{15}$N-leucine. A graph of the ppm errors for each identified ion is shown.[7] The mass of the peptide was increased by 1 Da in the C-terminal Trp derivative. **W** = modified Trp.

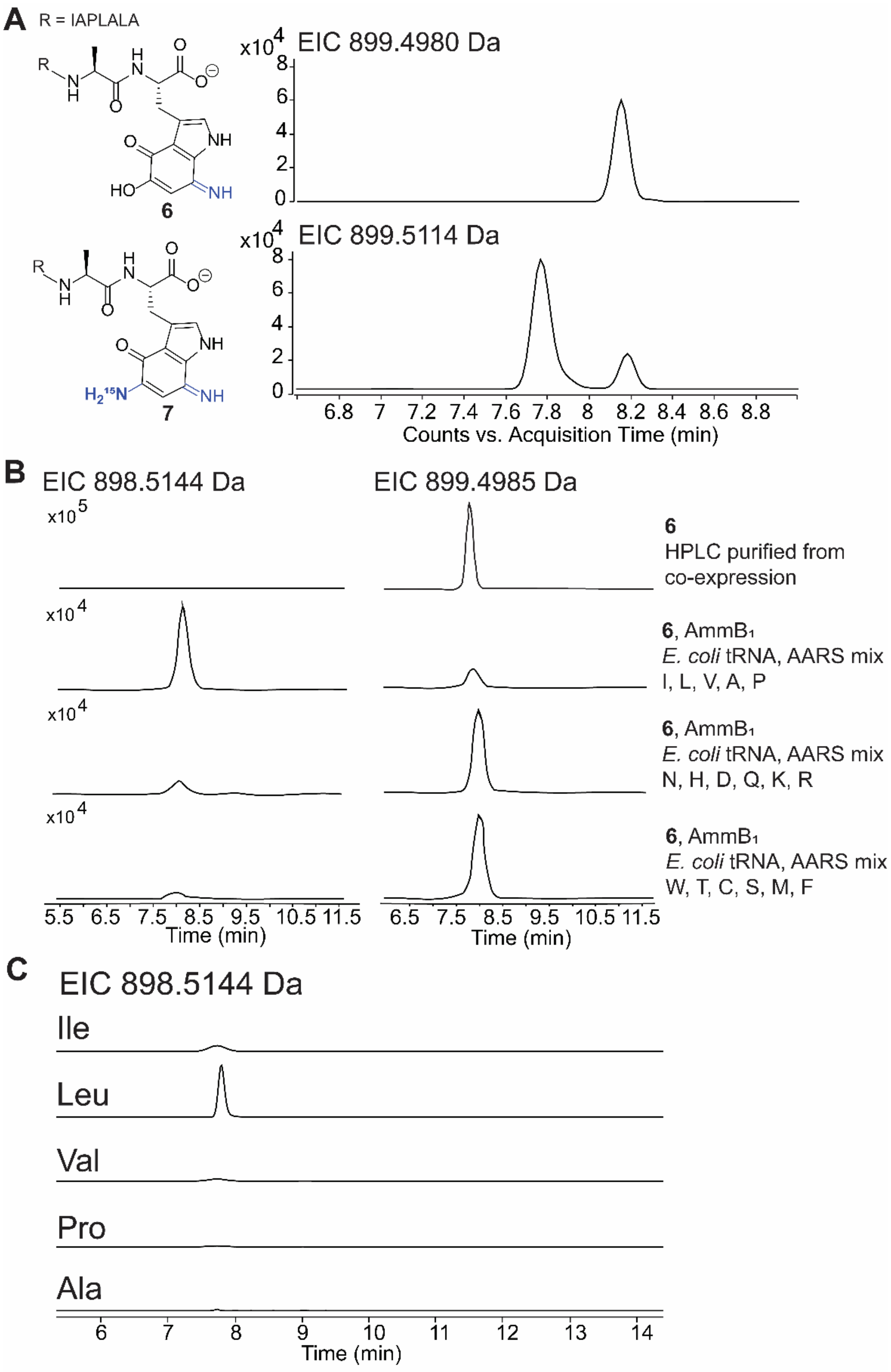


**Figure S14.** (**A**) LC-MS analysis of trypsin-digested aminoquinone intermediate **6** and the product **7** of the *in vitro* reaction with AmmB$_1$ using an $^{15}$N-labeled amino acid mix. The retention time of **7** is slightly earlier than that of **6**. Because of the use of $^{15}$N, the m/z of **6** and **7** is nearly the same in this experiment. (**B**) LC-MS analysis of *in vitro* reactions of **6** with *E. coli* tRNA and aminoacyl-tRNA synthetase (AARS) mix and the indicated mixture of amino acids. (**C**) LC-MS analysis of *in vitro* reactions of **6** with *E. coli* tRNA and AARS mix using individual Ile, Leu, Val, Pro, and Ala.

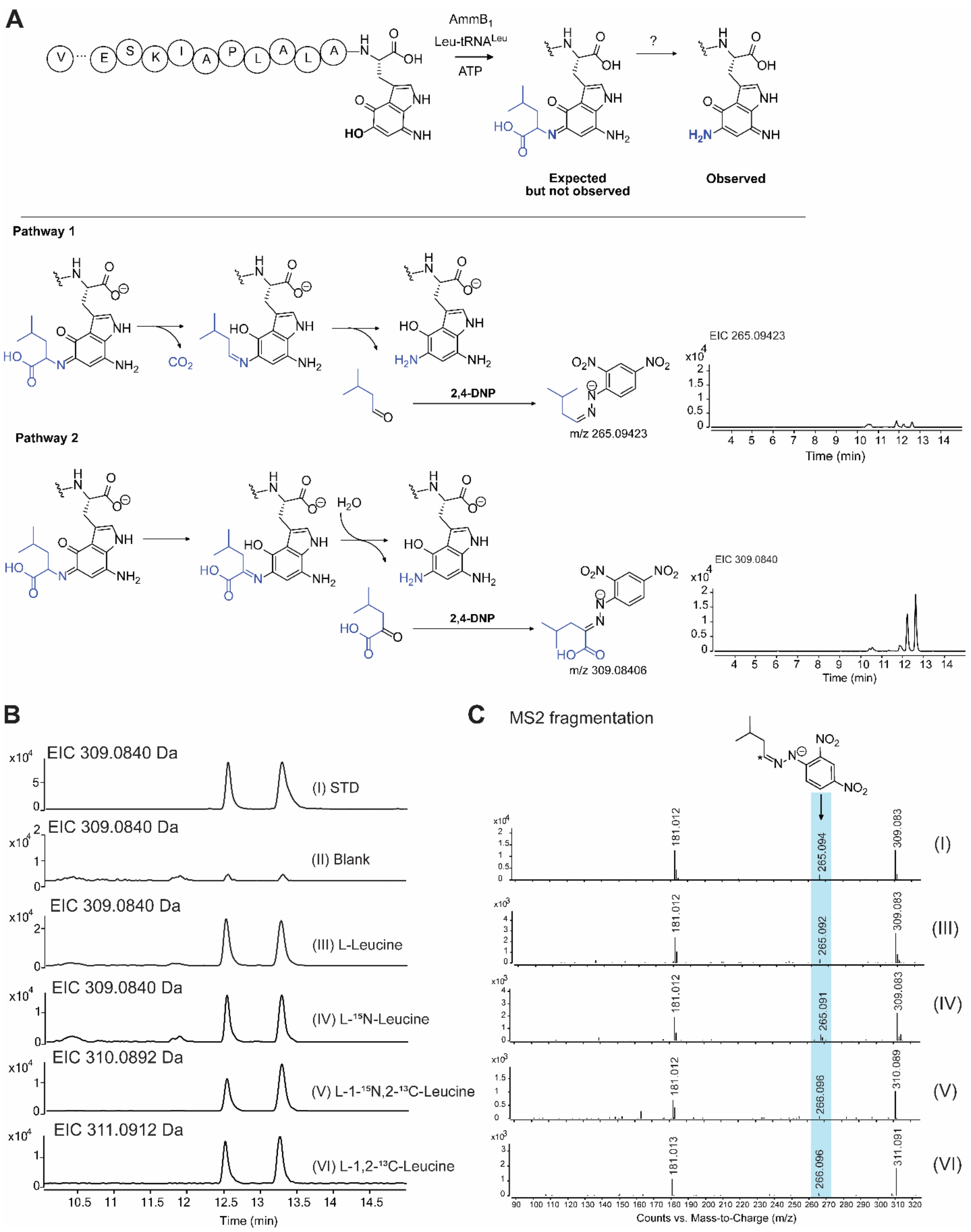


**Figure S15**. Phenylhydrazine derivatization of the AmmB$_1$ *in vitro* reaction. (A) Reaction product obtained in vitro by incubation of peptide **6** with AmmB$_1$, Leu, LeuRS, tRNA$^{Leu}$, and ATP. Two possible pathways are shown that explain the observed product. These mechanisms would generate the reduced form of the diamino product, consistent with the acetylated product generated in *E. coli* (Figure S13A). (B) EICs at the indicated m/z values of the top reaction performed with the isotopologs of Leu indicated in the graphs followed by derivatization with phenylhydrazine. Two stereoisomers (E/Z) are formed of the hydrazone. (C) The extracted mass spectra as well as the isotope distribution clearly show production of 4-methyl-2-oxovaleric acid.

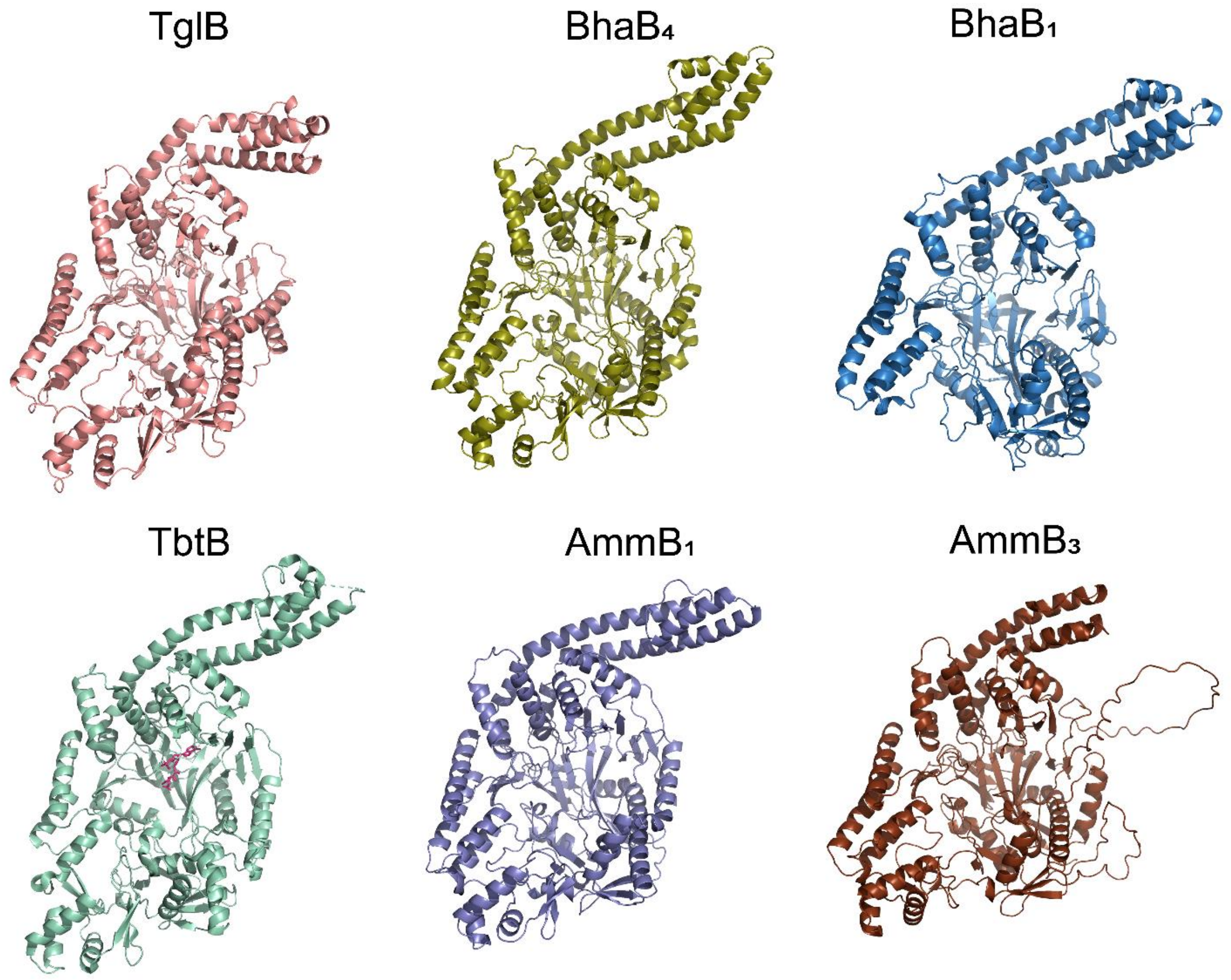


**Figure S16**. AlphaFold structures for the PEARLs TglB, $BhaB_4$, $BhaB_1$, $AmmB_1$, and $AmmB_3$ in comparison with the crystal structure of TbtB (PDB 6EC8).[8]

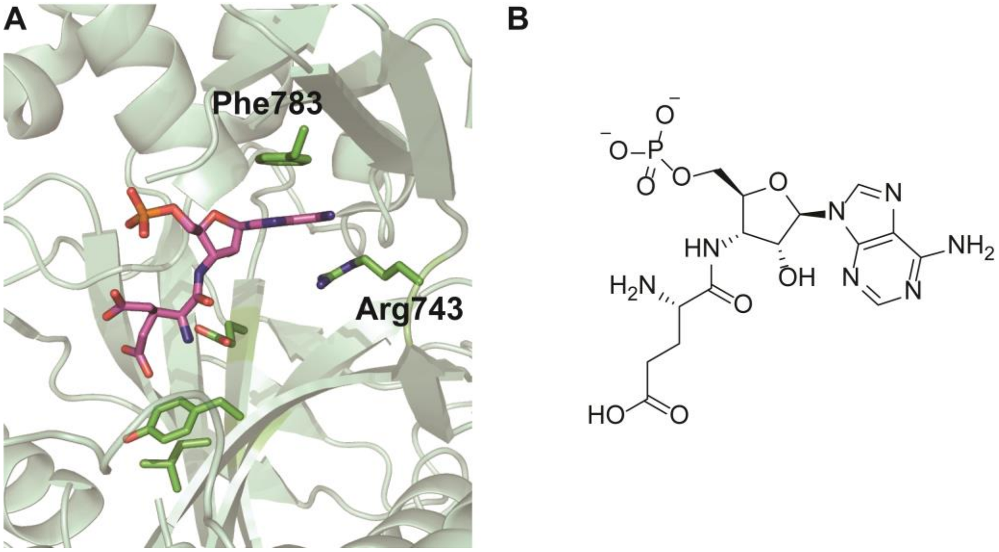


**Figure S17**. (A) Crystal structure of the split LanB glutamylation enzyme TbtB (PDB 6EC8). The active site is depicted with a nonreactive substrate analog, 5'-phosphoryl-desmethylglutamycin (B). Phe783 and Arg743 are positioned to provide π-π stacking and hydrogen bonding interactions with the adenosine moiety of the ligand.

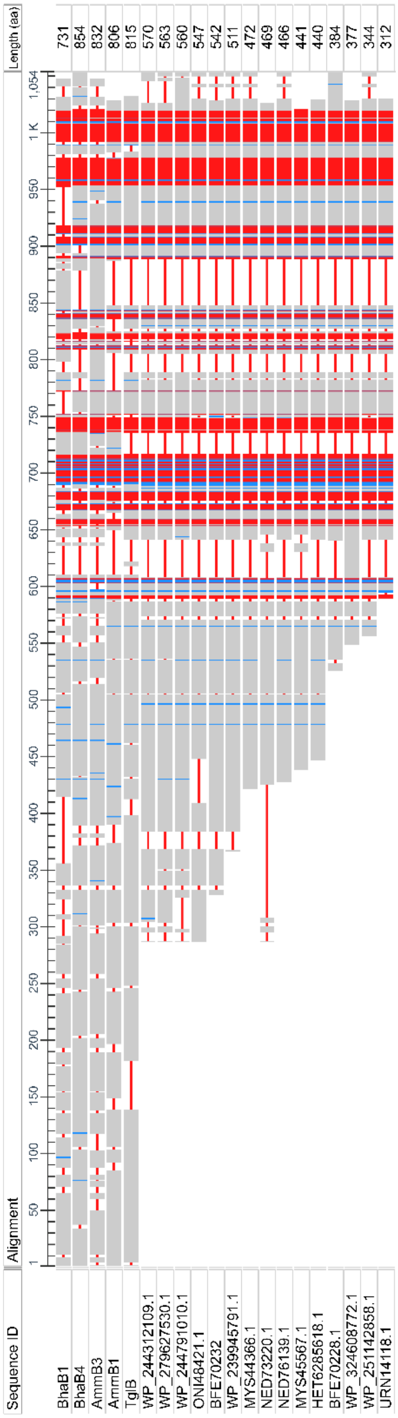


**Figure S18.** Constraint-based Multiple Alignment Tool (COBALT) was used to align the PEARLs $BhaB_1$, $BhaB_4$, $AmmB_1$, $AmmB_4$, and TglB versus shorter PEARL-like sequences found by a BlastP search with $AmmB_4$ as query. The red color indicates highly conserved regions and blue indicates less conserved regions. Shorter PEARL sequences align well to the C-terminus of the full length PEARL sequences. In addition, this region has structural homology with ATP-GRASP family members as seen from a 3D data-based search.

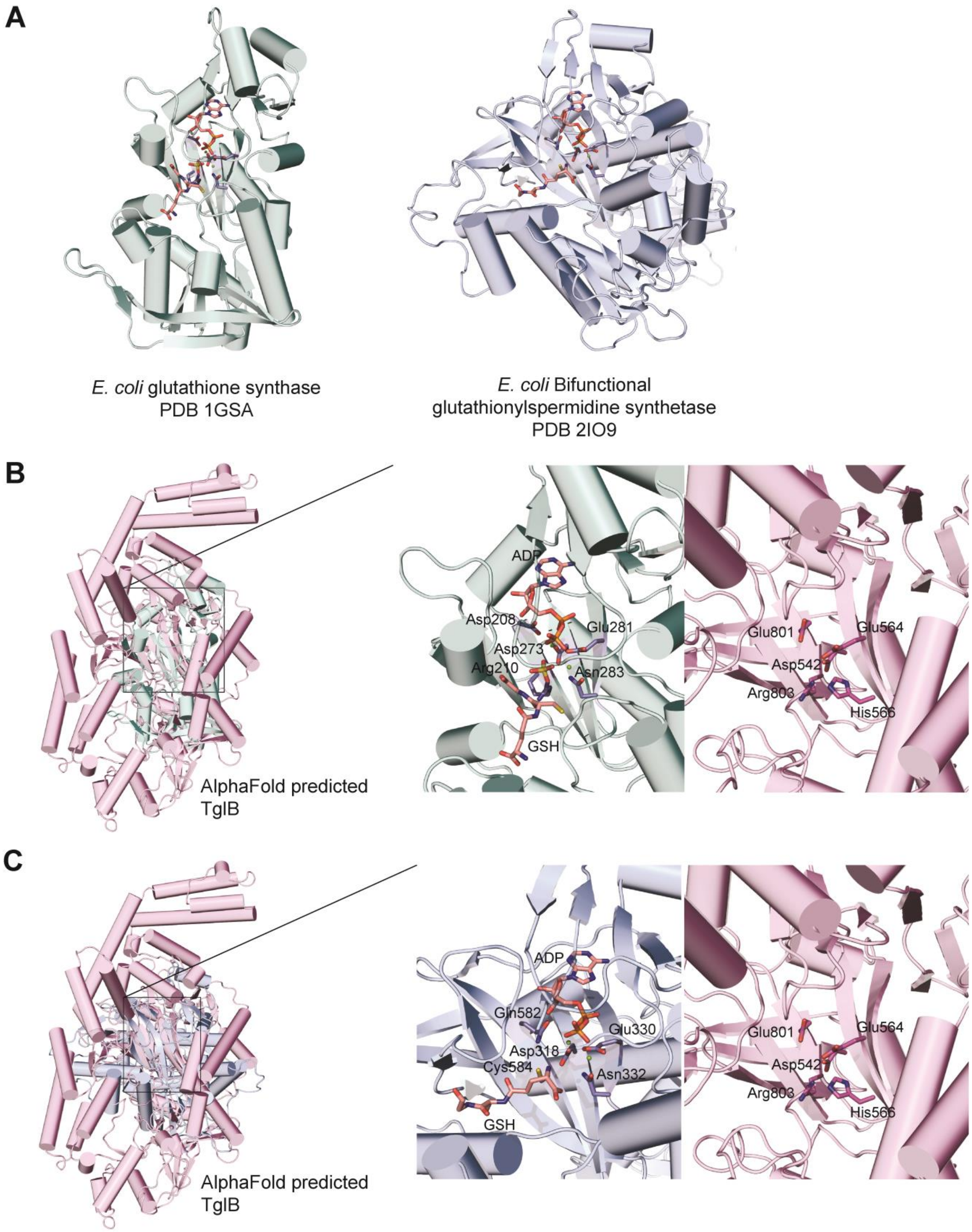
A
*E. coli* glutathione synthase
PDB 1GSA
*E. coli* Bifunctional
glutathionylspermidine synthetase
PDB 2IO9
B
AlphaFold predicted
TglB
ADP
Asp208
Glu281
Asp273
Arg210
Asn283
GSH
Glu801
Glu564
Asp542
Arg803
His566
C
AlphaFold predicted
TglB
ADP
Gln582
Glu330
Asp318
Cys584
Asn332
GSH
Glu801
Glu564
Asp542
Arg803
His566

**Figure S19.** (**A**) The AlphaFold predicted structure for TglB was overlayed with the crystal structure for *E. coli* glutathione synthase (PDB 1GSA) showing the potential binding site of ATP in the predicted PEARL structure. (**B**) Similarly, AlphaFold predicted structure for TglB was overlayed with the crystal structure for *E. coli* bifunctional glutathionylspermidine synthase (PDB 2IO9).

#### References

(1) Zhang, Z.; van der Donk, W. A. Nonribosomal peptide extension by a peptide amino-acyl tRNA ligase. *J. Am. Chem. Soc.* **2019**, *141*, 19625-33.
(2) Daniels, P. N.; Lee, H.; Splain, R. A.; Ting, C. P.; Zhu, L.; Zhao, X.; Moore, B. S.; van der Donk, W. A. A biosynthetic pathway to aromatic amines that uses glycyl-tRNA as nitrogen donor. *Nat. Chem.* **2022**, *14*, 71-7.
(3) Vaisar, T.; Urban, J. Probing the proline effect in CID of protonated peptides. *J. Mass Spectrom.* **1996**, *31*, 1185-7.
(4) Harrison, A. G.; Young, A. B. Fragmentation reactions of deprotonated peptides containing proline. The proline effect. *J. Mass Spectrom.* **2005**, *40*, 1173-86.
(5) Hayakawa, S.; Hashimoto, M.; Matsubara, H.; Turecek, F. Dissecting the proline effect: dissociations of proline radicals formed by electron transfer to protonated Pro-Gly and Gly-Pro dipeptides in the gas phase. *J. Am. Chem. Soc.* **2007**, *129*, 7936-49.
(6) Bleiholder, C.; Suhai, S.; Harrison, A. G.; Paizs, B. Towards understanding the tandem mass spectra of protonated oligopeptides. 2: The proline effect in collision-induced dissociation of protonated Ala-Ala-Xxx-Pro-Ala (Xxx = Ala, Ser, Leu, Val, Phe, and Trp). *J. Am. Soc. Mass Spectrom.* **2011**, *22*, 1032-9.
(7) Brademan, D. R.; Riley, N. M.; Kwiecien, N. W.; Coon, J. J. Interactive peptide spectral annotator: A versatile web-based tool for proteomic applications. *Mol. Cell. Proteom.* **2019**, *18*, S193-S201.
(8) Bothwell, I. R.; Cogan, D. P.; Kim, T.; Reinhardt, C. J.; van der Donk, W. A.; Nair, S. K. Characterization of glutamyl-tRNA-dependent dehydratases using nonreactive substrate mimics. *Proc. Natl. Acad. Sci. U. S. A.* **2019**, *116*, 17245-50.
